# Casein Kinase 1 dynamics underlie the PER2 circadian phosphoswitch

**DOI:** 10.1101/734624

**Authors:** Jonathan M. Philpott, Rajesh Narasimamurthy, Clarisse G. Ricci, Alfred M. Freeberg, Sabrina R. Hunt, Lauren E. Yee, Rebecca S. Pelofsky, Sarvind Tripathi, David M. Virshup, Carrie L. Partch

**Affiliations:** Department of Chemistry and Biochemistry, University of California Santa Cruz, Santa Cruz, CA 95064; Program in Cancer and Stem Cell Biology, Duke-NUS Medical School, Singapore 169857; Department of Chemistry and Biochemistry, University of California San Diego, La Jolla, CA 92093; Department of Pediatrics, Duke University Medical Center, Durham, NC 27710; Center for Circadian Biology, University of California San Diego, La Jolla, CA 92093

**Author notes:** Equal contributions (Jonathan M. Philpott, Rajesh Narasimamurthy, Clarisse G. Ricci).

**Keywords:** Circadian rhythms, Casein Kinase 1, phosphoswitch, activation loop, substrate recognition, non-consensus, conformational switch, Gaussian accelerated molecular dynamics, allostery, anion binding

## Abstract

Post-translational control of PERIOD stability by Casein Kinase 1δ and ε (CK1) plays a key regulatory role in metazoan circadian rhythms. Despite the deep evolutionary conservation of CK1 in eukaryotes, little is known about its regulation and the factors that influence substrate selectivity on functionally antagonistic sites in PERIOD that directly control circadian period. Here we describe a molecular switch involving a highly conserved anion binding site in CK1. This switch controls conformation of the activation loop to define substrate selectivity on mammalian PER2, thereby directly regulating its stability. Integrated experimental and computational studies shed light on the allosteric linkage between two anion binding sites that dynamically regulate kinase activity. We show that period-altering kinase mutations from humans to *Drosophila* differentially modulate this activation loop switch to elicit predictable changes in PER2 stability, providing a foundation to understand and further manipulate CK1 regulation of circadian rhythms.

## Introduction

Circadian rhythms are generated by a set of interlocked transcription/translation feedback loops that elicit daily oscillations in gene expression to confer temporal regulation to behavior, metabolism, DNA repair and more (Bass and Lazar, 2016). The PERIOD proteins (PER1 and PER2) nucleate assembly of large, multimeric complexes with the circadian repressors CRY1 and CRY2 that directly bind to and inhibit the core circadian transcription factor, CLOCK:BMAL1, on a daily basis (Aryal et al., 2017; Michael et al., 2017; Xu et al., 2015). PERs are stoichiometrically limiting for the assembly of these essential repressive complexes (Lee et al., 2011b). In this way, their abundance and post-translational modification state relay important biochemical information on the relative timing of the clock to other core clock proteins. Therefore, the expression, modification, and protein stability of PER1 and PER2 is under particularly tight regulation.

While both transcriptional and post-transcriptional mechanisms feature importantly in the rhythmic generation of PER proteins (Kojima et al., 2011; Takahashi, 2017), much attention has been focused on the post-translational control of PER stability orchestrated by its cognate kinases, CK1δ and the closely related paralog CK1ε (Hirano et al., 2016). These clock-associated kinases are somewhat unusual, in that they remain stably anchored to PER1 and PER2 throughout the circadian cycle (Aryal et al., 2017; Lee et al., 2001) via a conserved Casein Kinase Binding Domain (CKBD) (Eide et al., 2005; Lee et al., 2004). Mutations in CK1δ/ε (hereafter referred to jointly as CK1), as well as PER2, exert powerful control over circadian period, altering the intrinsic timing of circadian rhythms by hours *in vivo* (Lowrey et al., 2000; Toh et al., 2001; Xu et al., 2005; Xu et al., 2007). Because circadian period is linked to the timing of sleep onset, PER2 or CK1-dependent alterations to human circadian period manifest as sleep phase disorders that influence behavior and wellbeing on a daily basis (Jones et al., 2013).

PER2 is regulated by a CK1-dependent phosphoswitch, where kinase activity at two antagonistic sites functionally interact to control PER2 stability (Zhou et al., 2015). Two features define the CK1 phosphoswitch: degradation is initiated by phosphorylation of a Degron located several hundred residues upstream of the CKBD to recruit the E3 ubiquitin ligase, β-TrCP (Eide et al., 2005; Vanselow et al., 2006); this is counteracted by sequential phosphorylation of five serines embedded within the CBKD known as the FASP region (Narasimamurthy et al., 2018). This region is named for a Ser to Gly polymorphism in human PER2 that disrupts this stabilizing multi-site phosphorylation, shortens circadian period, and leads to Familial Advanced Sleep Phase Syndrome (Toh et al., 2001). Mutation of the Degron phosphorylation site has the opposite effect, stabilizing PER2 to significantly compromise circadian rhythms (Reischl et al., 2007) in a manner similar to its constitutive overexpression (Chen et al., 2009). Therefore, the balance of stabilizing and degrading phosphorylation by CK1 leads to a complex temporal pattern of degradation in PER2 that is important for circadian timing (Zhou et al., 2015).

Despite the importance of CK1 for circadian timing in eukaryotic organisms from humans to *Drosophila, Neurospora*, and green algae (Gorl et al., 2001; Kloss et al., 1998; van Ooijen et al., 2013; Xu et al., 2005), little is known about how its activity is regulated on clock protein substrates. CK1 is thought of as an anion- or phosphate-directed kinase, using negative charge on the substrate to template activity, for example, on a pSxxS consensus motif (Flotow et al., 1990). Biochemical studies exploring CK1 activity have primarily relied on the non-physiological acidic substrates casein (Venerando et al., 2014) and phosvitin (Lowrey et al., 2000), or used peptides harboring anion- or phosphate-driven motifs (Isojima et al., 2009; Marin et al., 1994; Shinohara et al., 2017). However, *in vitro* studies of clock-relevant kinase mutants using these non-physiological substrates have led to the puzzling conclusion that CK1 mutants that decrease or increase period length all have reduced kinase activity (Kivimae et al., 2008; Venkatesan et al., 2019).

This paradoxical observation motivated us to explore the molecular basis of CK1 activity on native PER2 substrates both *in vitro* and in cellular assays. To do this, we leveraged a comparative approach, examining multiple circadian mutants with a combination of cell-based, *in vitro*, structural and molecular dynamics methods. We discovered that the CK1 *tau* mutant (R178C) has reduced activity on the non-consensus priming of the FASP region as well as the downstream consensus sites, but exhibits a gain of function on the Degron site both *in vitro* and in cells (Gallego et al., 2006). Therefore, *tau* inverts PER2 substrate selectivity relative to the wild-type kinase to promote degradation of PER2. A mechanism for inverted substrate selectivity was suggested by the crystal structure of the *tau* kinase domain, which demonstrated the presence of a two-state conformational switch in the activation loop. Anion binding at a pocket near the activation loop biases the switch towards one conformation that correlates with a preference for the FASP substrate, while mutations that disfavor anion binding favor the alternate conformation with enhanced activity towards the Degron. Molecular dynamics studies reveal the conformation of the activation loop switch correlates with the substrate selectivity profile of WT and *tau* kinase. Specifically, we find the alternate conformation of the activation loop is stabilized in *tau*, forming the basis for its enhanced activity on the Degron. A comprehensive analysis of other short period kinase mutants from *Drosophila* to humans finds that they too differentially bias this intrinsic switch in substrate selectivity to enhance phosphorylation of the Degron and turnover of PER2. The anion-triggered activation loop switch may be a general mechanism regulating CK1 substrate selection.

## Results

### The *tau* mutant has decreased activity on the FASP region

The phosphoswitch model is predicated on CK1-dependent degradation of PER2 by phosphorylation of a β-TrCP-specific Degron upstream that is inhibited by stabilizing phosphorylation of the FASP region (Figure 1A). We recently demonstrated that CK1 primes FASP phosphorylation in a slow, rate-limiting step at serine 659 (mouse PER2 numbering, Figure 1B), with phosphorylation of the downstream consensus sites following rapidly in a sequential manner (dashed arrows at pSxxS consensus, Figure 1B) (Narasimamurthy et al., 2018). The *tau* mutation (R178C) was originally identified as a missense mutation that spontaneously arose in Syrian hamster CK1ε (Lowrey et al., 2000; Ralph and Menaker, 1988). Although *tau* enhanced Degron phosphorylation in cells to preferentially degrade PER2 (Gallego et al., 2006), the prevailing model was that a loss of function on the FASP region allowed for unchecked phosphorylation of the Degron. The mutation eliminates a positively charged residue in the first of three CK1 family-specific pockets (Sites 1, 2, and 3) that coordinate anions in the crystallographic structure (Figure 1C) (Longenecker et al., 1996). The Site 1 pocket where R178 sits is located adjacent to the active site and has been postulated to bind the phosphorylated priming site to position serines of the consensus motif at the active site (Longenecker et al., 1996; Zeringo and Bellizzi, 2014). Based on this model, it is predicted that *tau* should preferentially disrupt phosphorylation of the downstream consensus sites in the FASP region due to its inability to recruit the primed substrate.

**Figure 1.**
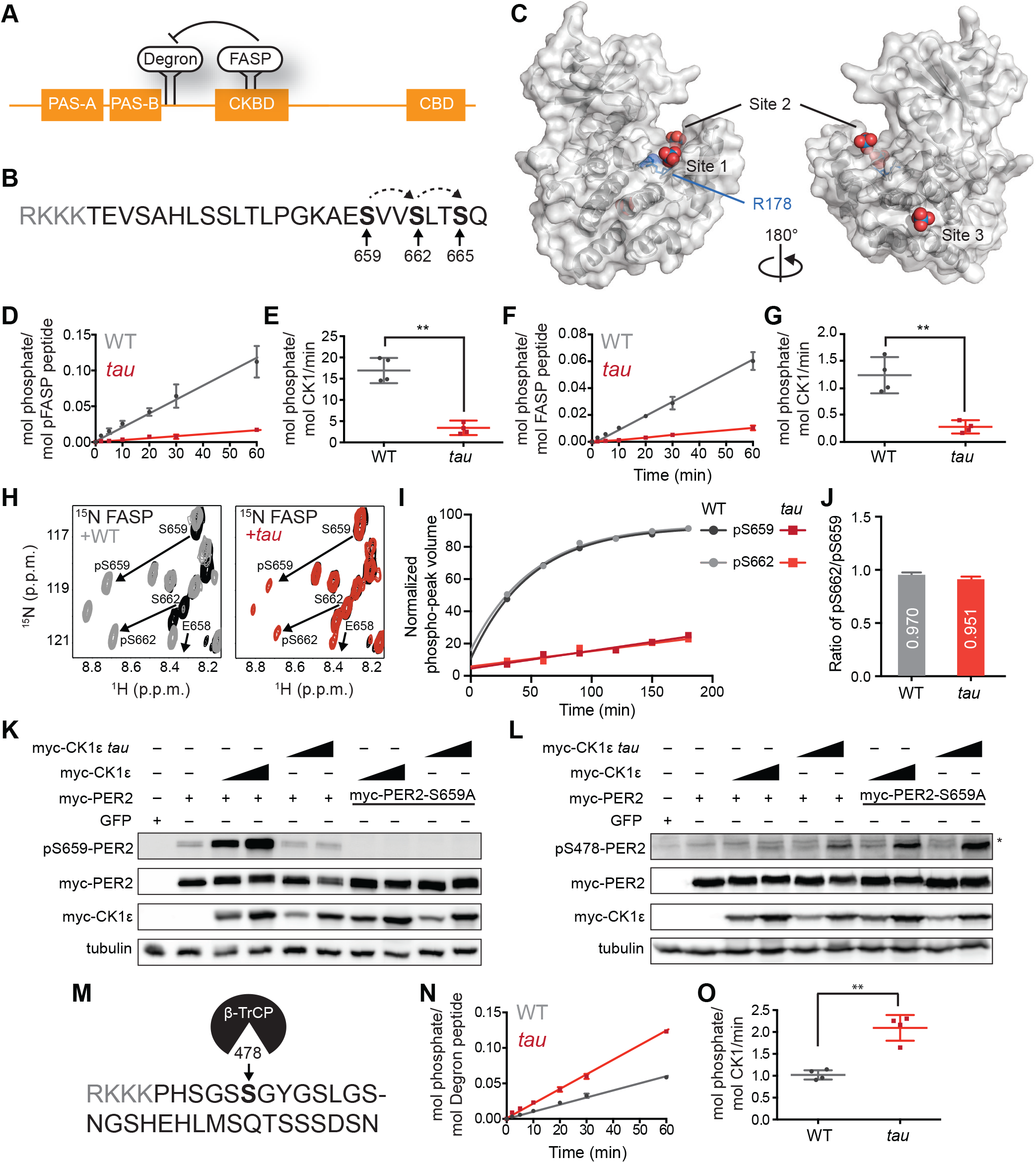
*tau* alters CK1 substrate selectivity on PER2 to enhance Degron phosphorylation. A, Domain map of PER2 with tandem PAS domains, Casein Kinase-binding domain (CKBD), CRY-binding domain (CBD) and CK1 phosphorylation sites. B, Sequence of the mouse PER2 FASP peptide with the priming site (S659, bold) and two downstream phosphorylation sites (S662 and S665, bold) that are phosphorylated sequentially by CK1δ (dashed arrows). Gray, polybasic motif included for ^32^P kinase assay. C, CK1δ kinase domain with 3 anion binding sites (PDB: 1CKJ with WO_4_^2-^ anions). R178, blue. D, Kinase assay with 20 nM CK1δ ΔC WT or *tau* on 200 μM of primed FASP peptide (pS659) (n = 4 with s.d.). E, Phosphorylation rates on primed FASP (n = 4 with s.d.). Significance assessed by unpaired Student’s two-sided t-test: **, p < 0.01. F, Kinase assay as in D, but with 200 nM CK1δ ΔC WT or *tau* on 200 μM of unprimed FASP (n = 4 with s.d.). G, Phosphorylation rates on the unprimed FASP peptide (n = 4 with s.d.). Significance assessed as above. H, Overlaid ^15^N/^1^H HSQC spectra at 3 hr timepoint in the NMR kinase assay on 200 μM ^15^N FASP (black) ± 1 μM WT (gray) or *tau* (red) CK1δ ΔC. Arrows, phospho-specific peaks corresponding to pS659 and pS662. I, Phosphoserine peak intensities for pS659 and pS662 by WT and *tau* kinases from NMR kinase assay. J, Ratio of consensus to priming activity on the FASP (pS662/pS659) in the NMR kinase assay. Errors were estimated from the standard deviation of the noise in the spectrum. K-L, Western blot of FASP priming site, detecting pS659 (K) or the Degron, detecting pS478 (L) on mouse myc-PER2 in HEK293 cell lysates after transfection with indicated expression plasmids. Representative blot from n = 3 shown. Wedge, 10 or 50 ng of myc-CK1ε plasmid used. *, non-specific band. M, Sequence of mouse PER2 Degron peptide with S478 (bold) and polybasic motif (gray). N, Kinase assay with 200 nM kinase on 200 μM Degron peptide (n = 4 with s.d.). O, Phosphorylation rates on Degron (n = 4 with s.d.). Significance assessed as above. See also Figure S1.

To test this idea, we used a FASP peptide based on the native mouse PER2 sequence (Figure 1B) that was primed synthetically by phosphorylation at S659. We also used a constitutively active version of the isolated wild-type (WT) or *tau* (R178C) kinase domain lacking its autoinhibitory tail (CK1δ ΔC, with 97% identity between CK1δ and CK1ε in the kinase domain). As expected, *tau* had significantly lower activity than the WT kinase on this primed substrate (Figure 1D-E). This was also true for the minimal, primed synthetic substrate CK1tide (Figure S1) (Shinohara et al., 2017). We then asked if *tau* could influence priming phosphorylation using an unmodified FASP peptide. As we observed before with WT kinase, phosphorylation of the non-consensus priming site occurs with much slower kinetics than the downstream consensus sites (Figure 1F-G) (Narasimamurthy et al., 2018). To our surprise, we found that *tau* also had significantly diminished activity on an unprimed FASP substrate (Figure 1F-G), indicating that R178 is also important for the non-consensus priming event. This decrease in activity of *tau* on the priming site was also validated using an ELISA-based kinase assay with an antibody that is specific for phosphorylated S659 (Figure S1).

To confirm that *tau* influences both priming and downstream events at the FASP region, we used a real-time NMR-based kinase assay. In contrast to the traditional kinase assays above, this assay provides site-specific information about modification of the substrate by measuring the increasing volume of new peaks that arise for phosphorylated serines over time (Theillet et al., 2013). Because we already established the dependency of priming to initiate sequential phosphorylation of downstream sites in the FASP region by CK1 (Narasimamurthy et al., 2018), this assay should differentiate the effects of *tau* on non-consensus priming and phosphorylation of the downstream consensus motif. If *tau* was simply deficient in recruitment of primed substrate, we should observe a similar degree of phosphorylation at the priming site (pS659) compared to WT kinase, but a decreased peak volume for the downstream serine (pS662). By contrast, we observed that the peaks for both pS659 and pS662 were both decreased in volume for *tau* (Figure 1H-I), with quantitative analysis of peak intensities revealing that the ratio of consensus to priming activity (pS662/pS659) was similar in both WT CK1 and *tau* (Figure 1J).

To test if these findings held in the context of full-length protein, we expressed myc-PER2 and WT or *tau* myc-CK1ε in HEK293 cells and assessed phosphorylation using an anti-pS659-specific antibody. Consistent with our *in vitro* kinase assays, the activity of *tau* was much lower on the FASP priming site relative to the WT kinase (Figure 1K). As expected, phosphorylation of the subsequent serine was also decreased with *tau* (Figure S1). To examine the possibility that loss of activity on the FASP region was specifically due to the missense cysteine mutation in the original mutant (R178C), we also tested an R178A mutant in the cell-based assay and found that it also exhibited much lower activity on the FASP priming site in a cell-based assay (Figure S1). Collectively, these data show that *tau* is deficient in both the slow priming step as well as the downstream sequential, primed phosphorylation of the FASP region.

### *tau* exhibits a gain of function on the Degron

Our previous model predicted that loss of function of *tau* for the FASP site allowed for unopposed phosphorylation of the Degron (Gallego et al., 2006). Using an antibody specific for phosphorylation of the CK1-dependent β-TrCP recruitment site at S478, we recapitulated the increased activity of *tau* on the Degron observed previously in myc-PER2 that was transiently expressed with CK1 in HEK293 cells (Figure 1L) (Gallego et al., 2006). Consistent with the phosphoswitch model that FASP phosphorylation antagonizes CK1 activity on the Degron (Figure 1A), we observed an increase in activity of WT kinase on the Degron with the S659A mutant that abrogates priming and downstream phosphorylation of the FASP region (Figure 1L) (Narasimamurthy et al., 2018). We also found that *tau* had increased activity on the Degron in the S659A mutant, demonstrating that the *tau* mutant can still be regulated by the phosphoswitch. Given that kinase activity on the Degron is clearly linked to phosphorylation of the FASP in cells, we sought to clarify whether *tau* truly exhibits increased activity at the Degron using a peptide-based kinase assay *in vitro* (Figure 1M). Here, we found that *tau* had significantly increased activity on the Degron relative to WT kinase (Figure 1N-O). These data suggest that in addition to any regulation imparted by the phospho-FASP on Degron activity in the context of full-length PER2, the *tau* mutation leads to a fundamental change in CK1 activity and substrate specificity.

### The *tau* mutation disrupts anion binding at Site 1 and Site 2 on CK1

To explore the molecular basis for *tau’s* altered substrate specificity, we solved a crystal structure of the CK1δ R178C kinase domain (Figure 2A and Table S1). Both WT and *tau* coordinate an anion at Site 3 similarly (Figure S2), but we observed a loss of anion binding at Sites 1 and 2 in *tau*. We expected that loss of the positively charged residue at position R178 would disrupt anion binding at Site 1, although the mutation led to only minor structural changes in this anion binding pocket (Figure 2B). *tau* had an alternate conformation of the activation loop near the second anion binding pocket that clashes with binding of the anion (Figures 2C), leading to its ejection from the site. This alternate conformation initiates at G175, three residues upstream of the *tau* mutation (Figure 2C). A rotation of the backbone at G175 to a left-handed configuration dramatically alters the configuration of upstream residues to create a distinct conformation of the activation loop (Figures 2D and S2). A backbone flip of a glycine at this conserved position has been observed in other serine/threonine kinases, linking changes in conformation of the activation loop to regulation of kinase activity (Nolen et al., 2004); therefore, the ‘loop up conformation observed in *tau* may lead to different kinase activity than the ‘loop down’ conformation observed in the WT kinase.

**Figure 2.**
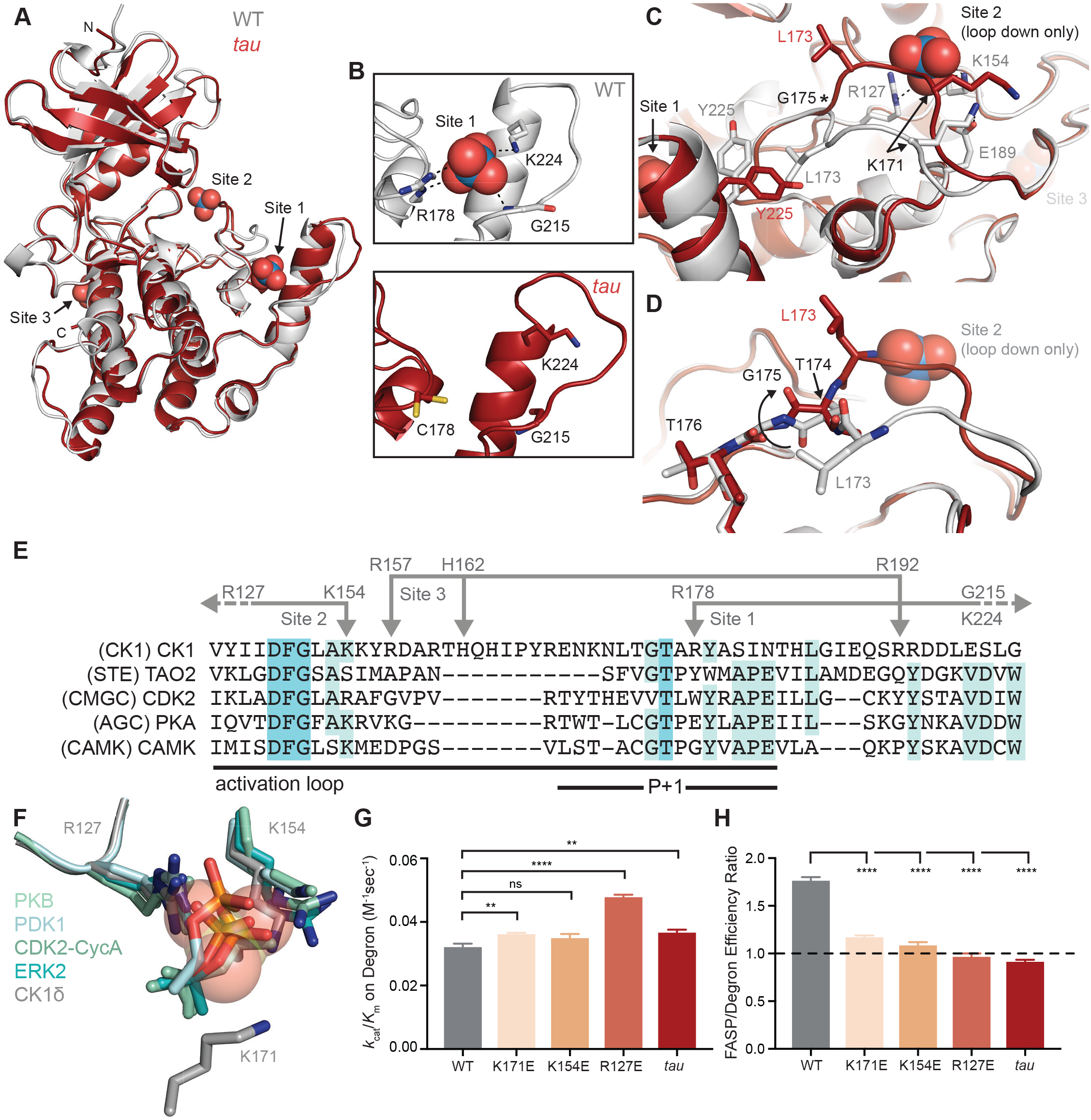
*tau* alters anion binding on CK1δ. A, Overlay of WT kinase domain (gray, PDB: 1CKJ, chain B) with *tau* (maroon, PDB: 6PXN, chain A). The 3 anion binding sites (WO_4_^2-^, from 1CKJ) are labeled. B, View of Site 1 in WT (top, gray) and *tau* (bottom, maroon). Polar interactions, dashed black lines. C, Overlaid view of Site 2 in WT and *tau* as above. Polar interactions, dashed black lines. Asterisk, hinge point for conformational change at G175. Note: Site 2 anion is only bound in WT, as it is blocked by the activation loop in *tau*. D, Representation depicting the left-hand configuration of G175 and subsequent rotation (solid arrow) of upstream residues T174 and L173. E, Alignment of the activation loop of CK1δ with representatives of other serine/threonine kinase families. Residues that coordinate anion binding on CK1 are indicated above in gray. F, Superposition of the Site 2 anion binding site in CK1 with the binding site for the phosphorylated activation loop of other serine/threonine kinases. Depicted are: PKB (PDB: 1O6K, pale cyan), PDK1 (1H1W, aquamarine), CDK2 (1QMZ, green cyan), ERK2 (2ERK, teal) and CK1δ (5×17, dark gray). Residues that coordinate the anion are depicted in sticks, as are phosphoserine or phosphothreonine residues from other kinases; the SO_4_^2-^ coordinated at Site 2 by CK1δ (PDB: 5×17) is shown in transparent spheres. G, Enzymatic efficiency on the Degron (n = 3 with s.d.). Significance assessed relative to WT with an unpaired Student’s two-sided t-test: **, p < 0.01; ***, p < 0.001; ****, p < 0.0001. H, Ratio of enzymatic efficiency on FASP relative to Degron (n = 3 with s.d.). Equivalent activity on FASP and Degron, dashed line. Significance assessed as above. See also Figure S2.

The activation loop is the key feature that distinguishes the CK1 family from other serine/threonine kinases. A deviation in sequence from the highly conserved ‘APE’ motif in the P+1 region, which defines specificity for the residue that follows the phosphotarget, is largely attributed as the reason for a lack of mechanistic insight into CK1 substrate selectivity (Figure 2E) (Goldsmith et al., 2007). The activation loop and surrounding region also contain residues that coordinate the three anions observed in nearly all CK1 structures: the residues that coordinate binding at Sites 1 and 3 are unique to the CK1 family, while R127 and K154, corresponding to Site 2, are broadly conserved in other kinase families. R127 is part of the highly conserved HRD motif that is used by some kinases to coordinate a phosphorylated residue in the activation loop and regulate substrate binding (Figure 2F) (Johnson et al., 1996). CK1 family kinases are generally considered to be constitutively active because they do not require phosphorylation of the activation loop (Goldsmith et al., 2007). However, the activity of clock-relevant kinases CK1δ and CK1ε is inhibited by autophosphorylation of their disordered C-terminal tails (Graves and Roach, 1995; Rivers et al., 1998) and potentially by the phosphorylated FASP region (Figure 1L). Therefore, the anion binding sites could represent the basis for a CK1-specific regulatory mechanism by facilitating the binding of phosphorylated C-terminal tails or substrates, and/or anionic signaling molecules (Fustin et al., 2018; Kawakami et al., 2008).

### Eliminating anion binding at Site 2 differentially regulates CK1 activity on the FASP and Degron

Mutation of the residues corresponding to positions R127 and K154 at Site 2 severely decreases the activity of kinases that depend on phosphorylation of the activation loop (Gibbs and Zoller, 1991; Leon et al., 2001; Skamnaki et al., 1999). To test the role of Site 2 binding in regulating CK1 activity, we made charge reversion mutants at positions R127 and K154, as well as at K171 located nearby on the activation loop, and measured enzymatic efficiency (*k*_cat_/*K*_m_) on FASP and Degron peptides *in vitro*. We observed a modest decrease (~25%) in activity towards the FASP peptide (Figure S2 and Table S2), suggesting that the ‘loop down’ conformation that is enforced by anion binding at Site 2 may be important for FASP activity. Strikingly, the same mutations increased activity towards the Degron, with a ~50% gain in efficiency for R127E (Figure S2 and Figure 2G). To further examine the role of K171 in regulation of anion binding, we solved a structure of the K171E mutant crystallized in high sulfate conditions and found a full complement of three anions bound (Figure S2). In our structure, the N170 sidechain replaced the mutant E171 to make an interaction with the sulfate, demonstrating that local flexibility in the activation loop allows it to retain sulfate binding at Site 2 to some degree. The kinetic data suggest that variability in the strength of anion binding at the Site 2 pocket correlates with kinase activity on the FASP and Degron peptides. Notably, a decrease in activity on the stabilizing FASP region and an increase in activity on the Degron makes the Site 2 coordination mutants much more *tau*-like (Figure 2H). This is consistent with a recent report that both *tau* and charge reversion mutants at Site 2 lead to decreases in PER2::LUC stability in cell-based assays (Shinohara et al., 2017).

### The activation loop switch is intrinsic to the CK1 family of kinases

In looking at both copies of the kinase found in the asymmetric unit of the *tau* crystal, we discovered that the mutant kinase can take on either the ‘loop down’ or the ‘loop up’ conformation (Figure 3A). The activation loop is not stabilized by crystal contacts in the alternate ‘loop up’ conformation and was explicitly modeled based on good density in both conformations (Figure S3), suggesting that the kinase has an intrinsic ability to take on two discrete conformations in a switch-like manner. Since the first structure of CK1δ published over two decades ago, most have been determined after crystallization with high concentrations of sulfate or citrate anions (Long et al., 2012; Longenecker et al., 1996; Minzel et al., 2018). Because anion binding at Site 2 is incompatible with the activation loop in its ‘loop up’ conformation, prior crystallographic conditions have disfavored this alternate conformation. The WT structure that we used for our analysis (PDB: 1CKJ) was first crystallized with a low concentration of anions, and then derivatized with tungstate as an analog for phosphate before data collection (Longenecker et al., 1996). Importantly, this structure also displays the same two discrete conformations marked by translocation of residue L173 (Figure 3A), confirming that the activation loop switch is an intrinsic property of the CK1δ kinase that has not been explored functionally.

**Figure 3.**
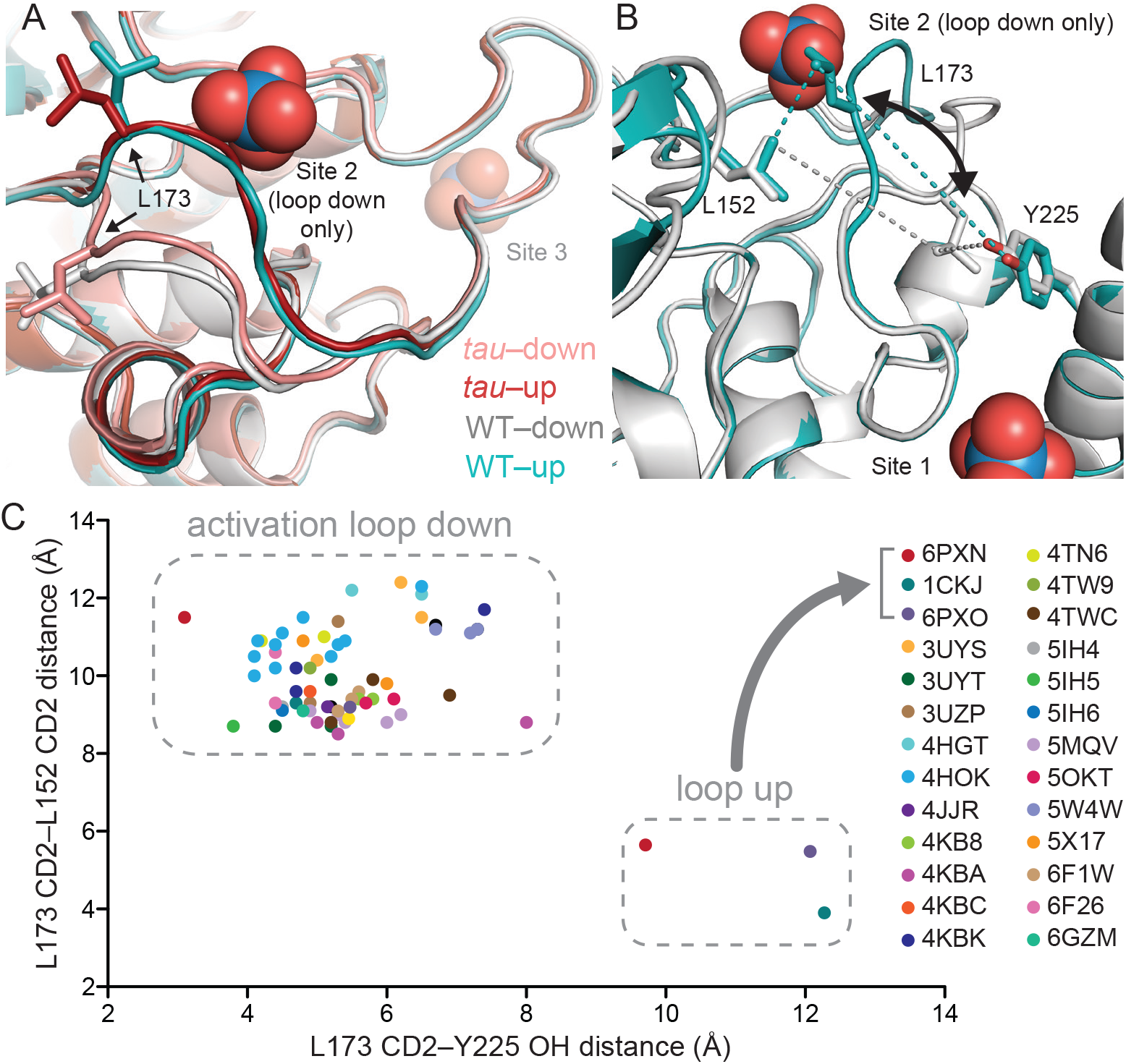
*tau* alters an intrinsic molecular switch in the activation loop of CK1δ. A, View of the activation loop switch in WT (PDB: 1CKJ, chains A (cyan) and B (gray)) and *tau* (PDB: 6PXN, chains A (maroon) and B (salmon)). B, The position of L173 CD2 relative to either L152 CD2 or Y225 OH reports on the conformation of the activation loop in the ‘loop down’ (gray) or ‘loop up’ (cyan) conformation in WT CK1δ (PDB: 1CKJ). C, Scatter plot of interatomic distances in Å (from panel B) measured in 68 chains from 26 different crystal structures of CK1δ ΔC. See Figure S3 and Supplemental Table 3 for more information.

We then probed the library of existing CK1δ structures using the positioning of L173 as a quantitative metric for activation loop conformation by measuring interatomic distances of the L173 sidechain CD2 atom to either CD2 of L152 (short distance in the ‘loop up’ conformation, long in the ‘loop down’) or the hydroxyl of Y225 (long in the ‘loop up’, short in the ‘loop down’) (Figure 3B). A survey of 68 chains from 26 different crystal structures of CK1δ (in at least 7 space groups) demonstrated that *tau* and the WT kinase from PDB entry 1CKJ are the only structures of CK1δ to have ever been captured in the ‘loop up’ conformation (Figure 3C). Moreover, the residues that coordinate anion binding are broadly conserved in the CK1 family (Figure S6), and resident anions are observed in structures of other CK1 family kinases (e.g., CK1ε, CK1γ3; Table S3), implicating anion binding and its regulation of the activation loop in a mechanism that may be generally conserved in the CK1 family. To determine if we could independently capture the two states of the activation loop switch in WT kinase, we optimized sulfate-free crystallographic conditions and solved the structure of WT CK1δ. Here, similar to the 1CKJ WT structure, we also found both conformations of the activation loop in the two molecules of the asymmetric unit (Figure S3). Therefore, both conformations of activation loop switch can occur at low anion concentrations in WT kinase, but the *tau* mutation seems like it might favor the ‘loop up’ conformation because it was observed in crystals that grew in the presence of high sulfate concentrations. Altogether, these data suggest that the *tau* mutation may allosterically regulate anion binding at Site 2 via the activation loop.

### *tau* stabilizes the rare ‘loop up’ conformation of the CK1 activation loop

To probe the dynamic behavior of WT CK1 and see how the enyzme is perturbed by the *tau* mutation, we performed Gaussian Accelerated Molecular Dynamics (GaMD) simulations (Miao et al., 2015) on four systems: WT and *tau* CK1 with the activation loop in the crystallographically-defined ‘up’ or ‘down’ conformations (Table S5). By monitoring the Root Mean Square Deviation (RMSD) of the activation loop with respect to the ‘down’ or ‘up’ crystallographic conformations, we set out to assess its stability over the course of 500 ns simulations (Figure 4A-D). We found that the activation loop remained stably in position when simulations were started from the ‘loop down’ conformation for both *tau* and WT. Similar results were seen when the anion was computationally removed from Site 2, suggesting that this conformation of the activation loop is intrinsically stable (Figure S4). However, in simulations starting from the ‘loop up’ conformation, the WT activation loop rapidly underwent a conformational change, as shown by increased RMSD^up^ values. Because we did not see a concomitant decrease in the RMSD_down_ values, we can conclude that this is not a complete transition from ‘loop up’ to ‘loop down’ on this timescale. Importantly, these transitions occurred more frequently in WT CK1 than in *tau* (Figure 4C-D). This confirms that the ‘loop up’ conformation is better tolerated in *tau*, consistent with our observation of this apparently rare conformation in our crystal structure (Figure 3).

**Figure 4.**
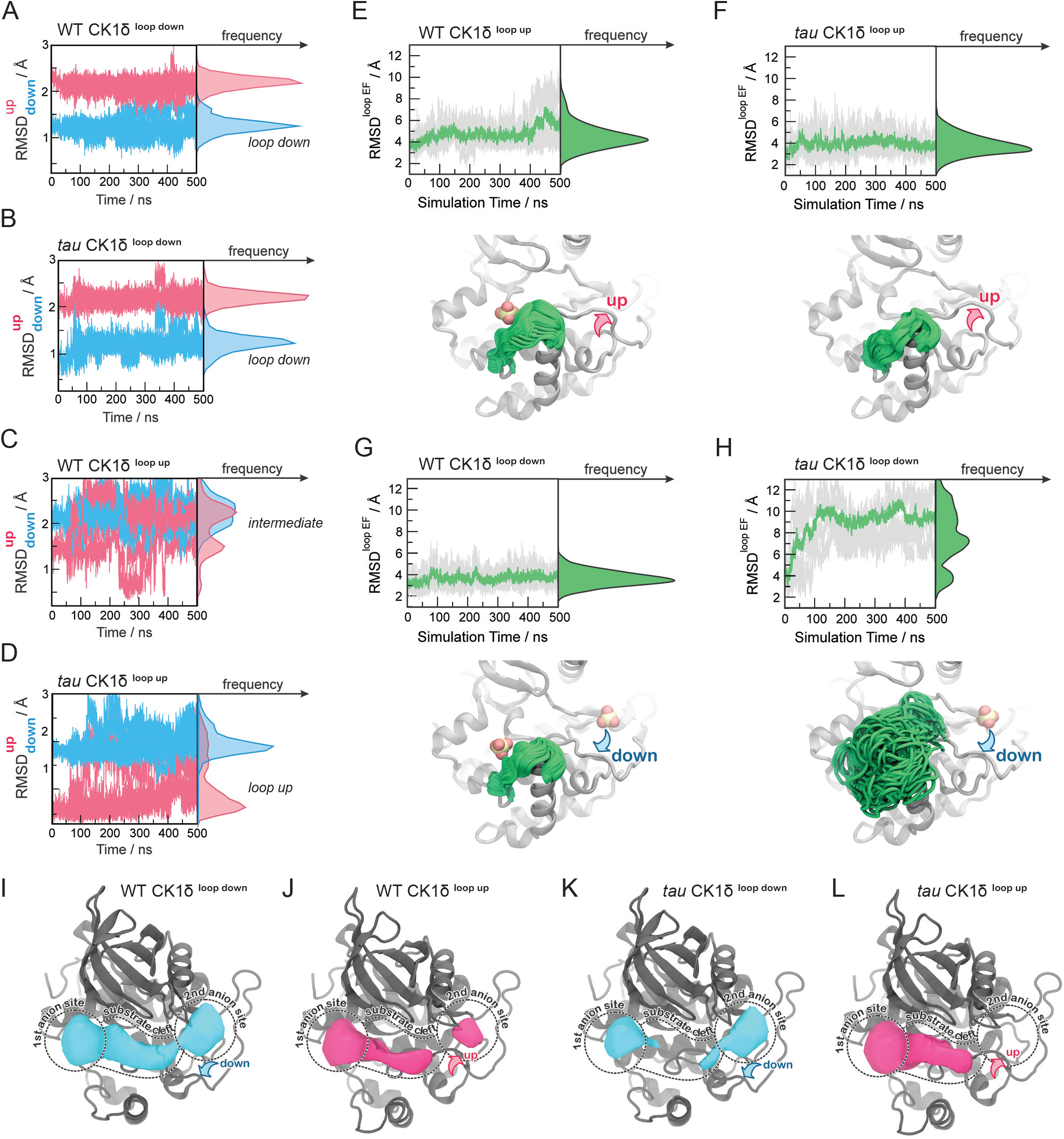
Probing the dynamics of CK1δ with GaMD simulations. A-D, Stability of the activation loop assessed by the RMSD of residues 168-175 with respect to the ‘loop down’ (RMSD_down_, blue) or ‘loop up’ conformation (RMSD^up^, pink), as observed in the crystal structure. For each system, the RMSDs from all five MD replicas are superimposed. Panel A, WT CK1δ^loop down^; B, *tau* CK1δ^loop down^; C, WT CK1δ^loop up^; D, *tau* CK1δ^loop up^. E-H, Dynamics of the EF loop assessed by the RMSD of residues 213-224 with respect to the initial structure. For each system, the RMSD was calculated for individual replica (gray lines, n = 5) and then averaged (green). The molecular representations in panels E-H show the crystallographic structure of CK1δ (gray) superimposed with snapshots of the L-EF loop extracted from the GaMD simulations (green). When present in the crystal structure and the simulation, sulfate anions are represented by spheres. I-L, Alterations in anion and substrate-binding clefts arise from the activation loop switch and *tau* mutation. Volumes for the binding clefts were extracted and averaged from GaMD simulations in the four states: panel I, WT CK1δ^loop down^; J, WT CK1δ^loop up^; K, *tau* CK1δ^loop down^; L, *tau* CK1δ^loop up^. Water and anions were removed from the analysis. Volumetric maps are contoured at 0.1 and represent regions that were consistently open during the simulations. See also Figure S4.

### The activation loop allosterically controls the dynamics of loop L-EF in *tau*

While monitoring the overall dynamics of WT and *tau* CK1, we detected a major difference in the dynamics of the loop connecting α-helices E and F (loop L-EF, Figure S4), which is part of the anion binding site disrupted by the *tau* mutation (Figure 4E-H). This is intriguing because temperature-dependent dynamics of loop L-EF were recently shown to be important for the temperature compensated activity of CK1 on PER2 (Shinohara et al., 2017). In our simulations of the WT kinase, this loop exhibited relatively restricted mobility, regardless of whether the activation loop was in the ‘up’ or ‘down’ conformation. Similar results were observed when the anion was computationally removed from Site 2 (Figure S4). However, we observed that the ‘loop down’ conformation of the activation loop in *tau* was accompanied by a strong disorganization of loop L-EF (Figure 4H), leading to significantly larger conformational freedom compared to the WT kinase (Figure 4G). Surprisingly, the enhanced dynamics of loop L-EF was not observed when *tau* was in the ‘loop up’ conformation; instead, loop L-EF displayed the same restricted dynamics as the WT kinase (Figure E-F). These data suggest that the disruption of Site 1 by loop L-EF dynamics in the *tau* mutation may be due to allosteric communication with the activation loop, and by proxy, anion binding at Site 2.

### *tau* dynamically reshapes the substrate-binding cleft in CK1

To investigate how the activation loop conformation and dynamics of loop L-EF might influence substrate selectivity, we calculated the volume of the substrate binding cleft and adjacent anion binding sites throughout the 500 ns GaMD simulations (Figure 4 and S4). As expected, Site 2 was only completely open only when the activation loop was in the ‘loop down’ conformation for the WT kinase (Figure 4I). Interestingly, this site became partially open in simulations of the WT kinase starting from the ‘loop up’ conformation (Figure 4J), indicating that the intermediate conformational state observed in our simulations might allow the kinase to recover, to some extent, the ability to bind an anion at Site 2. The conformational state of the activation loop indirectly affects the volume of the substrate binding cleft with opposing effects in WT and *tau*. In WT, the substrate binding cleft was open more consistently with the activation loop in its preferred ‘down’ conformation (Figure 4I-J). By contrast, the substrate binding cleft was open more consistently in *tau* in the ‘loop up’ conformation (Figure 4L) and, due to the dynamic disordering of loop L-EF, often closed when in the ‘loop down’ conformation (Figure 4K). Because the activation loop does not contact the substrate binding cleft, it cannot directly affect the shape of the cleft by steric effects. Instead, closing of the substrate binding cleft in *tau* occurs due to the conformational disorganization in loop L-EF, which is allosterically induced when the activation loop is in the ‘down’ conformation.

### *tau* influences the global dynamics of CK1

We next used principal component analysis to uncover effects of *tau* on the principal modes of motion displayed by CK1 during the GaMD simulations (Figure 5, S5 and Supplemental Movie 1). The 1^st^ principal component consisted of a clear ‘open-and-close’ movement of the enzyme, achieved mainly by dislocation of the N-terminal lobe (N-lobe) with respect to the top of the helix F (Figure 5A). This mode of motion has been shown to control accessibility to the ATP-binding site and regulate substrate access in other kinases (McClendon et al., 2014). The histograms of the 1^st^ principal component show that WT CK1 sampled more of the open conformations compared to *tau* (Figure 5C). The 2^nd^ principal component consisted of a twisting movement of the N-lobe with respect to the top of helix F and significant rearrangement of loop L-EF, which can either be extended or collapsed (Figure 5B). When collapsed, loop L-EF had the effect of sterically closing the substrate binding cleft. The histograms of the 2^nd^ PCA illustrate that loop L-EF adopts more extended conformations in WT CK1 and more collapsed conformations in *tau* (Figure 5D). The collapsed conformations require loop L-EF to undergo a significant conformational change, which agrees with the high conformational freedom seen in this region when the *tau* activation loop is in the ‘loop down’ conformation. Altogether, the GaMD simulations suggest that the *tau* kinase never behaves fully like the WT enzyme; compared to WT, it stabilizes the rare, Degron-preferring conformation of the activation loop that remodels the substrate binding cleft and excludes anion binding at Site 2. By contrast, when *tau* samples the ‘loop down’ conformation of the activation loop, it leads to a dynamic disordering of loop L-EF that favors conformations of the kinase that likely decrease its enzymatic efficiency on the FASP region.

**Figure 5.**
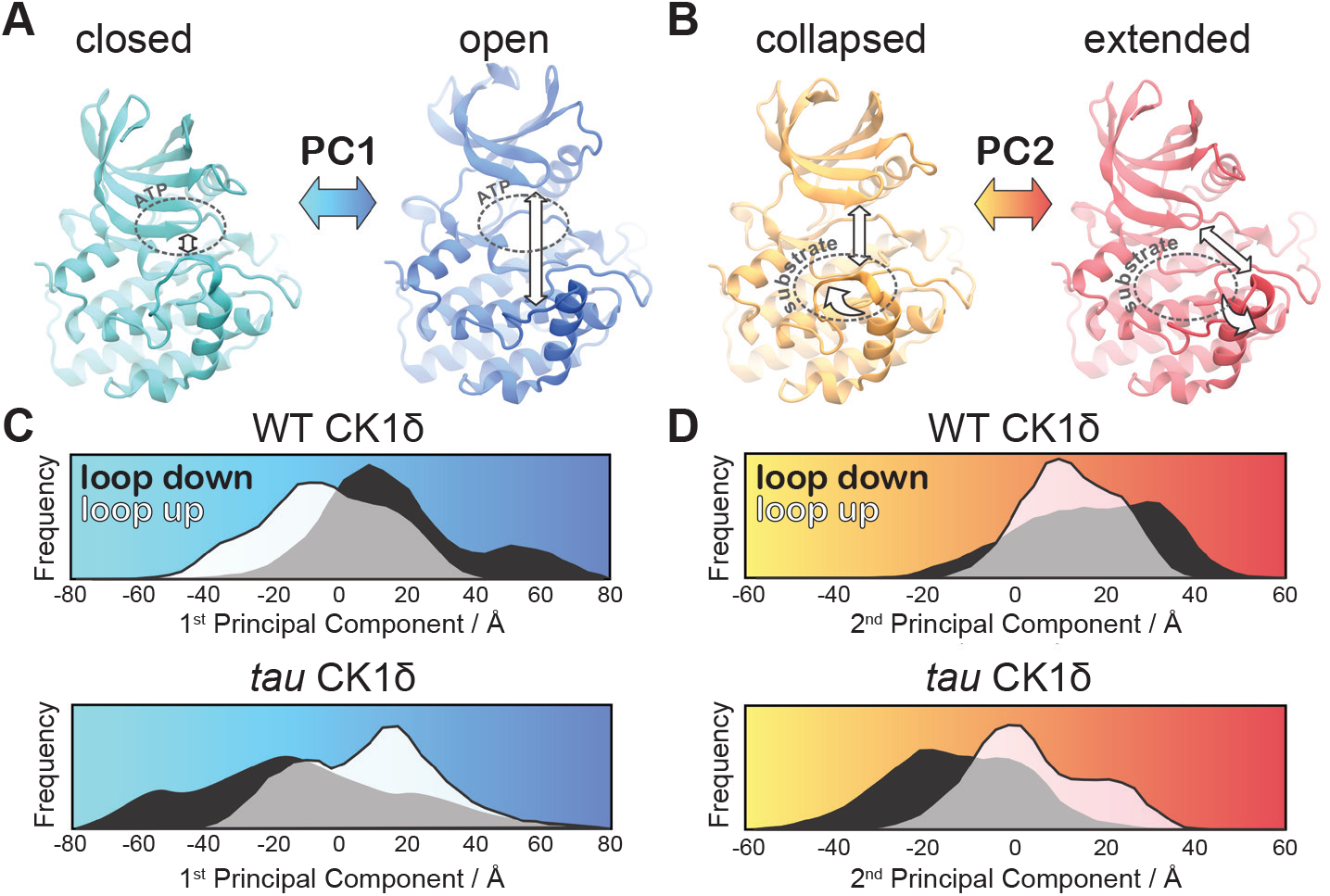
*tau* influences the principal modes of motion in CK1δ. Principal modes of motion obtained from Principal Component Analysis (PCA). A, PC1 consists of an ‘open-and-close’ movement achieved by dislocation of the N-lobe with respect to the top of the helix F to control accessibility of the ATP-binding site (dotted circle). B, PC2 consists of a twisting movement of N-lobe with respect to the top of helix F and significant rearrangement of the loop L-EF, which can be extended or collapsed against the substrate binding site (dotted circle). C-D, Structural representations correspond to (panel A) PC1 = −80 Å (cyan), PC1 = 80 Å (blue) and (panel B) PC2 = −60 Å (orange), and PC2 = 60 Å (red). The histograms represent projections of the accumulated GaMD trajectories along the 1^st^ (C) or 2^nd^ (D) principal components for WT CK1δ and the *tau* mutant, either in the activation ‘loop down’ (black) or ‘loop up’ (white) conformations. See also Figure S5 and Supplemental Movie 1.

### Circadian alleles from *Drosophila* to humans occur throughout CK1

We mapped known mutant alleles that influence circadian rhythms onto the CK1 structure to gain further insight into its mechanism of regulation (Figure 6). *Doubletime* (DBT), the CK1δ/ε ortholog in *Drosophila*, has one allele that causes a short circadian period (*dbt^S^*, P47S) while all others lead to a long period (Kloss et al., 1998; Rothenfluh et al., 2000; Suri et al., 2000). Many long period mutations occur at or near catalytically important residues. The classic *dbt^L^* allele M80I (Kloss et al., 1998) is directly adjacent to the catalytic lysine, K38; notably, expressing low levels of the catalytically dead K38R mutant in *Drosophila* also leads to a long period (Muskus et al., 2007). *dbt^AR^* and *dbt^G^* correspond to H126Y and R127H of the catalytically important HRD motif at Site 2 (Rothenfluh et al., 2000; Suri et al., 2000); together, these long period mutations sandwich the catalytic DFG motif and the regulatory spine that controls kinase activity (Figure 6B) (Taylor and Kornev, 2011). Two loss of function alleles occur in the activation loop: *dco^18^* (S181F) is located right behind the *tau* site (R178), linking it to the substrate binding channel and activation loop, while *dco^2^* (G175S) occurs at the hinge point for the activation loop switch (Zilian et al., 1999). Although *Drosophila* and mammalian PER proteins are somewhat functionally divergent, conservation of their Degrons (Chiu et al., 2008; Eide et al., 2005) and FASP-like stabilizing phosphorylation sites (Kivimae et al., 2008; Top et al., 2018; Xu et al., 2007) suggests that there may be some conservation in their regulation by CK1 (Figure 6C). In line with this, expressing mammalian CK1δ with the *tau* or *dbt^s^* mutation leads to short period circadian rhythms in flies (Fan et al., 2009). Moreover, there is an incredible degree of conservation in the CK1 family; the entire surface-exposed area linking Sites 1 and 2 and the substrate binding cleft are ≥95% identical in 20 species from humans to unicellular green alga where CK1 has been implicated in regulation of circadian clocks (Figure S6).

**Figure 6.**
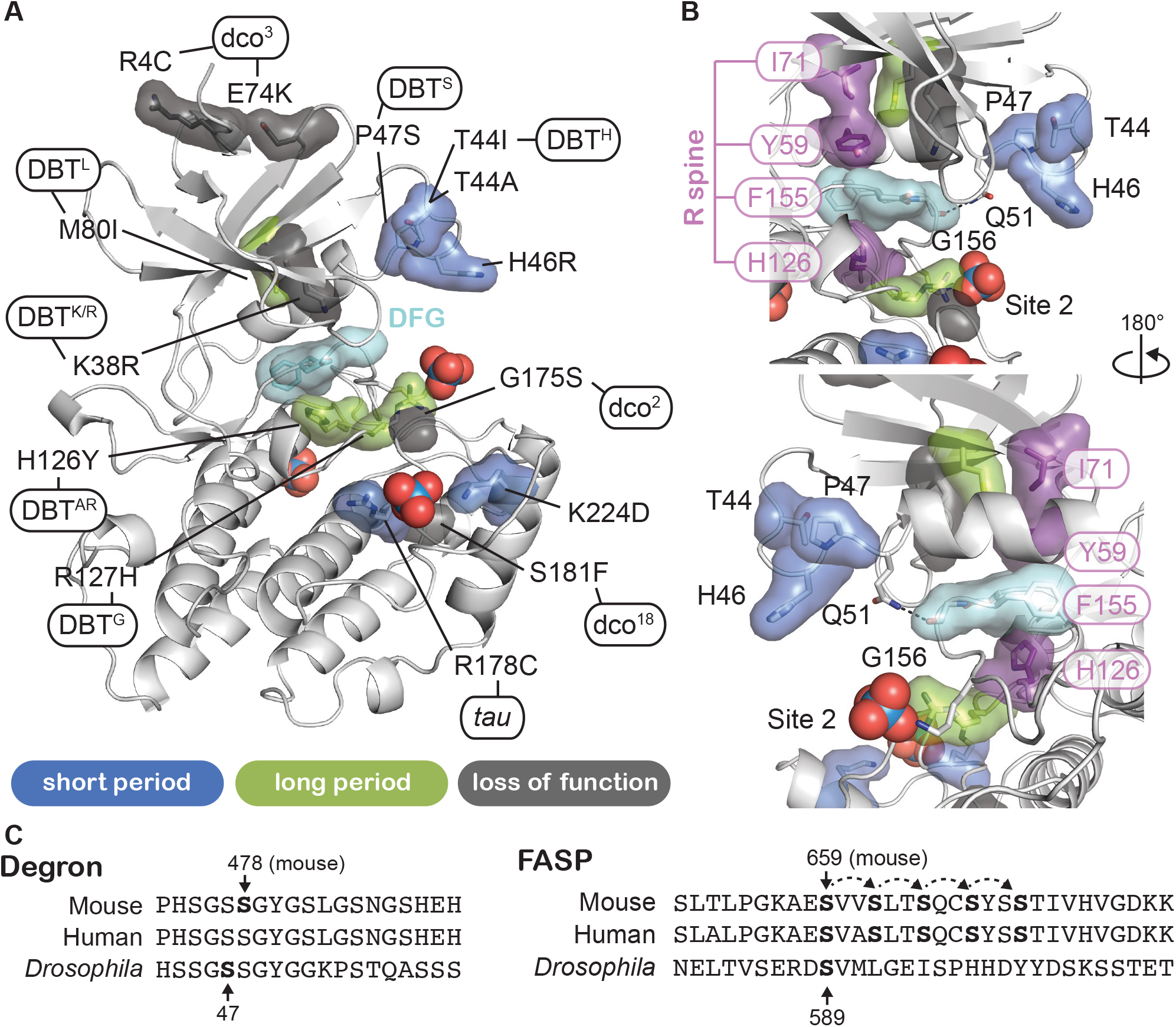
Proximity of CK1 alleles map to catalytic and substrate binding sites. A, Structure of CK1δ (PDB: 1CKJ, chain B) clock relevant alleles mapped from mammalian CK1 or *Drosophila* DBT. Mutants are colored by phenotype: short period (blue), long period (green), loss of function (gray). DFG catalytic motif, cyan. B, View of N-lobe period mutants and the regulatory spine (R-spine, purple) with F155 of the DFG motif in cyan. Polar interactions between Q51 and G156 that link the N- and C-lobe are depicted with a dashed black line. C, Alignment of the mammalian and *Drosophila* Degron and FASP/stabilizing sequences. Residues in bold have experimental support for phosphorylation. Dashed arrows indicate sequential phosphorylation following the consensus pSxxS motif.

### Other short period mutants exhibit differential activity on the FASP and Degron

We were intrigued by three short period mutants from humans and *Drosophila* (T44A, H46R, and P47S) that colocalize in the N-lobe right above Site 2 (Figure 6B). Given the changes in N-to C-lobe dynamics that we observed in our simulations of CK1 (Figure 5), we wondered how these mutations would influence substrate selectivity. We first tested the activity of these short period kinase mutants in cell-based transfection assays by monitoring both FASP priming (Figure 7A) and Degron (Figure 7B) phosphorylation. Similar to the Site 2 mutants we tested earlier (Figure S2), the short period mutants T44A, H46R, and P47S each retained substantially more kinase activity on the FASP priming site than *tau*. A triple mutant of all three short period mutants (3M) did not have additive effects. Likewise, they also appeared to retain kinase activity or even exhibited modest increases in Degron phosphorylation relative to WT (Figure 7B), although not to the same degree as *tau*. On balance, it appears that the Site 2 mutants (short period or anion binding) might act differently from *tau* in that they retain FASP priming activity while still increasing activity at the Degron relative to WT.

**Figure 7.**
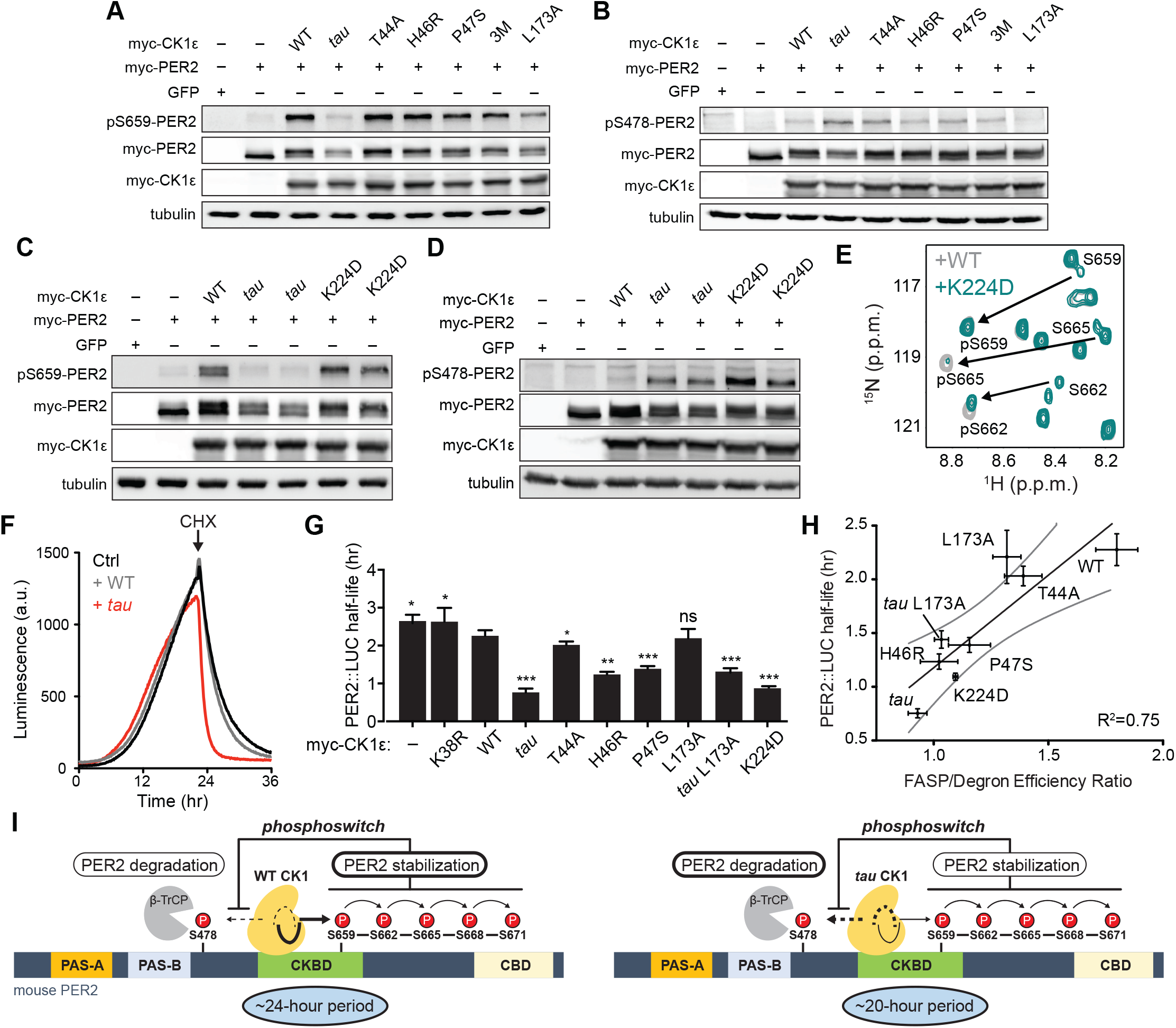
Substrate discrimination on the PER2 phosphoswitch is regulated by the CK1 activation loop switch. A,B Western blot of FASP priming site, detecting pS659 (A) or Degron site, detecting pS478 (B) phosphorylation on mouse myc-PER2 in HEK293 cell lysates after transfection with indicated myc-CK1ε expression plasmids. 3M triple mutant: T44A, H46R, P47S. Representative blot from n = 3 shown. C,D Western blot of FASP priming (C) or Degron (D) phosphorylation on PER2 as above with the myc-CK1ε K224D mutant. Representative blot from n = 3 shown with replicate samples loaded for *tau* and K224D. E, Overlaid ^15^N/^1^H HSQC spectra at 3 hr timepoint in the NMR kinase assay on 200 μM ^15^N FASP + 1 μM K224D (teal) or WT (gray) CK1δ ΔC. Arrows, phosphospecific peaks corresponding to pS659, pS662, and pS665. F, Representative real-time luminescence data for PER2::LUC stability in HEK293 cells transfected with myc-PER2::LUC plus empty vector (black) or myc-CK1ε WT (gray) or *tau* (red) as indicated (n = 4). 40 μg/mL cycloheximide (CHX) added 24 hours post-transfection (arrow). G, Quantification of PER2::LUC half-life with different myc-CK1ε mutants. Data represent mean ± s.d. (n = 4) with significance assessed as above. H, Scatterplot with linear regression analysis of the ratio of enzyme efficiencies (*k*_cat_/*K*_m_) for FASP and Degron relative to the PER2::LUC half-life determined in panel G and Figure S7. All data are plotted as mean ± s.d. (n = 4 for PER2::LUC and n = 3-4 for enzyme efficiencies). Black, linear regression to data; gray, 95% confidence interval. I, The conformational switch of the CK1δ/ε activation loop is coupled to substrate selection in the PER2 phosphoswitch. Left, the activation loop of the WT kinase is stable in the ‘loop down’ conformation, leading to preferential phosphorylation the FASP region, which stabilizes PER2 by reducing phosphorylation of the Degron. Right, the activation loop of *tau* kinase is better tolerated in the alternate ‘loop up’ conformation leading to an intrinsic gain of kinase function on the Degron and loss of kinase function on the stabilizing FASP region. This switch in substrate preference promotes PER2 degradation and leads to a shorter circadian period. CKBD, CK1 binding domain; CBD, CRY binding domain. See also Figure S7.

We also examined another short period mutant located at Site 1, K224D, which was recently reported to have a ~20-hour circadian period (Shinohara et al., 2017). Like the other short period mutants positioned near Site 2, K224D retained FASP priming activity (Figure 7C). However, in contrast to them, K224D exhibited a sharp increase in Degron phosphorylation in HEK293 cells relative to WT that was similar to *tau* (Figure 7D). This effect was not dependent on the charge inversion, as we saw the same effect with a K224A mutant (Figure S7). Notably, our *in vitro* assessment of K224D enzyme efficiency showed an overall decrease in efficiency on the FASP peptide containing multiple serines (Table S2). Taken together, these data suggest that priming of the FASP site S659 is intact, but subsequent downstream phosphorylation might be compromised in the mutant. To test this, we used the NMR-based kinase assay to provide single residue resolution on the stepwise phosphorylation of the FASP region (Narasimamurthy et al., 2018). Consistent with the cellular data, K224D retained normal priming activity at S659, but phosphorylation of the downstream serines that conform to the pSxxS consensus motif was compromised relative to WT kinase (Figure 7E). Just like blocking phosphorylation of the FASP region with the S659A mutant (Figure 1L), loss of phosphorylation at these downstream sites could allow for the increased kinase activity on the Degron that we observed in cellular assays. Therefore, although *tau* and K224D both enhance Degron phosphorylation to a similar degree and lead to ~20-hour circadian periods (Lowrey et al., 2000; Shinohara et al., 2017), they likely achieve this through different mechanisms on the kinase—*tau* dynamically disrupts Site 1 in the FASP-preferring ‘loop down’ conformation of the activation loop to reduce both priming and downstream phosphorylation of the FASP region, while K224D (and K224A) likely just disrupt the anion binding at Site 1 that is important for phosphorylation of the pSxxS consensus motifs.

### The ratio of FASP/Degron enzyme efficiency correlates with PER2 stability

One commonality of short period mutants should be enhanced Degron phosphorylation in cells, as both cellular (Gallego et al., 2006; Vanselow et al., 2006) and *in vivo* studies of short period mutants demonstrate that PER2 stability is decreased in a CK1-dependent manner (Price et al., 1998; Xu et al., 2007). To test this model, we measured the effect of mutant CK1 co-expression on the half-life of PER2::LUC in real-time after cycloheximide treatment (Figure 7F and S7). All of the mutants, apart from L173A, lead to a shorter half-life of PER2::LUC (Figure 7G). We mutated L173 because it appeared to serve as a latch for the activation loop in both its ‘loop up’ and ‘loop down’ conformations (Figure 2C). We found that the mutant substantially decreased kinase activity at both the FASP priming and Degron sites in the transfection-based assay and *in vitro* (Figure 7A-B and Table S2), and a double mutant with *tau* decreased its activity on the Degron to a similar degree (Figure S7). We hypothesized that the L173A mutant might have a half-life similar to WT because its reduced phosphorylation of the stabilizing FASP region was offset by decreased activity on the Degron (Figure 7G and Table S2). This suggested to us that the ratio of intrinsic kinase activity on the stabilizing FASP region relative to the Degron might account for CK1’s ability to initiate PER2 degradation by β-TrCP (Eide et al., 2005; Vanselow et al., 2006). Indeed, Figure 7H shows that a significant correlation exists between PER2::LUC half-life and the intrinsic ratio of CK1 kinase activity on these two key sites that establish the phosphoswitch mechanism. Taken with the other aspects of this study, our data strongly suggest that CK1 uses activation loop dynamics to orchestrate substrate specificity and contribute directly to the phosphoswitch regulation of PER2 stability, and therefore, the timing of circadian rhythms (Figure 7I).

## Discussion

Despite its powerful control over the timing of circadian rhythms along the entire branch of eukaryotes from humans to green algae (van Ooijen et al., 2013; Xu et al., 2005), very little is known about the molecular determinants of CK1δ substrate selectivity and activity. Here we identify a conformational switch in the activation loop of CK1 that regulates its activity on two distinct non-consensus motifs that underlie the phosphoswitch controlling PER2 expression and circadian timing in mammals (Zhou et al., 2015). Anion binding at Site 2 and the intrinsic dynamics of the conformational switch bias the WT kinase towards phosphorylation of the FASP region to stabilize PER2. We show that reducing anion binding at Site 2 regulates the activation loop to reshape the substrate binding cleft and enhance activity on the Degron. Using a series of mutations in the kinase that shorten circadian period from *Drosophila* to humans, we demonstrated that all exhibit enhanced activity on S478 in the Degron, consistent with our earlier study (Gallego et al., 2006). However, only the *tau* mutant has decreased activity on the FASP priming site and downstream serines. Molecular simulations revealed that enhanced dynamics of loop L-EF likely underlie the decreased activity of the *tau* mutant CK1 on the FASP region. Moreover, they also demonstrated that the activation loop switch can exert allosteric control over the dynamics and conformation of loop L-EF through the *tau* site.

Allosteric regulation is a common feature of protein kinases, often based on dynamic changes in ensembles of residues that can occur in the absence of major changes in conformation (Kornev and Taylor, 2015). CK1 has remarkable, histone-like conservation (≥ 95% identity) of the entire surface-exposed area linking the two key anion binding sites and substrate binding cleft, suggesting that the mechanisms we discovered here likely apply broadly to circadian rhythms as well as other CK1-regulated processes in other eukaryotes. Binding of regulatory anions at these conserved sites could arise from phosphorylated CK1 itself in *cis* via its autoinhibitory tail (Rivers et al., 1998), or in *trans* from the binding of phosphorylated substrates like the FASP region, to allow for the generation of feedback regulation directly on the kinase. In line with this, changes in the sequence (Fustin et al., 2018) or phosphorylation (Eng et al., 2017) of the autoinhibitory tail of CK1δ/ε could alter the balance of FASP and Degron phosphorylation to control circadian period. In this way, our study demonstrates that the dynamics of CK1δ/ε directly encode its activity in the PER2 phosphoswitch (Zhou et al., 2015). The conformational equilibrium of the activation loop may also play a role in the temperature-compensated activity of CK1δ/ε observed *in vitro* that is linked to the dynamics of loop L-EF, an insertion in the clock-relevant kinases CK1δ and CK1ε that plays a key role in temperature compensation of circadian rhythms (Shinohara et al., 2017).

We used PER2 stability here as a cellular proxy to study the effect of mutations from CK1δ and CK1ε on circadian timing. PER proteins seem to have a special role in the mammalian clock as state variables that define both the timing and phase of circadian rhythms through changes in their abundance (Balsalobre et al., 2000; Zylka et al., 1998). CK1-dependent changes in PER abundance likely affect circadian period based on their role as stoichiometrically limiting factors in the assembly of transcriptional repressive complexes in the feedback loop of the molecular clock (Aryal et al., 2017; Kim and Forger, 2012; Lee et al., 2011b). Recent studies have shown that CK1 is maintained in an active state while bound to PER2 *in vitro* (Qin et al., 2015), which may allow it to target other clock proteins for phosphorylation in the repressive complexes (Aryal et al., 2017), a property that seems to be conserved in *Drosophila* (Yu et al., 2006). Therefore, more studies are needed to fully understand the interplay between PER2 stability, PER-CK1 interactions and the regulation of post-translational modifications, including by phosphatases (Lee et al., 2011a) and other kinases (Hayasaka et al., 2017; Hirota and Kay, 2009; Oshima et al., 2019), that ultimately control circadian rhythms.

CK1 is highly active on primed (pSxxS) sites (Narasimamurthy et al., 2018), supporting its designation as the canonical consensus motif of the kinase. However, we find it compelling that many of the biologically important roles of CK1δ/ε and the related kinase CK1α as key regulators of Wnt signaling (Marin et al., 2003), the DNA damage response (Knippschild et al., 1997), cell cycle (Penas et al., 2015), and circadian rhythms (Kloss et al., 1998; Lowrey et al., 2000) depend on their activity on non-consensus sites. The ability of CK1δ/ε to phosphorylate these lower affinity, nonconsensus sites on PER2 is likely dependent on the formation of a stable, stoichiometric complex with PER2 via two highly conserved sites that flank the FASP phosphorylation region in the CKBD (Eide et al., 2005; Lee et al., 2004). The generally low activity of CK1 that we observe on clock-relevant non-consensus sequences may also be important for the slow timescale of circadian rhythms. This property is also conserved in KaiC, the enzyme that controls circadian timing in the cyanobacterial system (Abe et al., 2015), suggesting commonalities in the biochemical origins of building slow biological clocks.

## Acknowledgments

We would like to thank Danny Forger, Jae Kyoung Kim, Yinglong Miao, and J. Andrew McCammon for useful discussions and Sivakumar Parthiban for technical assistance. We also thank the beamline staff for their assistance at the Advanced Photon Source beamline 23-ID-D and Advanced Light Source beamline 8.3.1, as well as the San Diego Supercomputer Center (SDSC) for technical support. This work was funded by the National Medical Research Council of Singapore Grant NMRC/CIRG/1465/2017 (to D.M.V.) and National Institutes of Health Grants R01 GM031749 (to C.G.R.), GM107069 and R01 GM121507 (to C.L.P.), as well as funds from the NIH Office of the Director under Award S10 OD018455 for the 800 MHz NMR spectrometer used here. S. Hunt was supported by NRSA F32 GM133149.

## Author contributions

Conceptualization, J.M.P., R.N., C.G.R., D.M.V., and C.L.P.; Methodology, Investigation, J.M.P., R.N., C.G.R., A.M.F., S.R.H., L.Y., and R.P.; Validation, J.M.P., R.N., S.T.; Formal Analysis, J.M.P., R.N., C.G.R.; Visualization, J.M.P., R.N., C.G.R., and C.L.P.; Writing–Original Draft, J.M.P., C.G.R., and C.L.P.; Writing–Review and Editing, J.M.P., R.N., C.G.R., A.M.F, S.R.H., D.M.V., and C.L.P.; Supervision, D.M.V., and C.L.P.; Funding Acquisition, D.M.V., and C.L.P. All authors consented on the final draft of the manuscript.

