## Supplemental Figures and Materials for "Casein Kinase 1 dynamics underlie the PER2 circadian phosphoswitch"

This file includes:
Supplemental Figures 1-7
Supplemental Tables 1-5
Supplemental Movie 1
Materials and Methods
Supplemental References

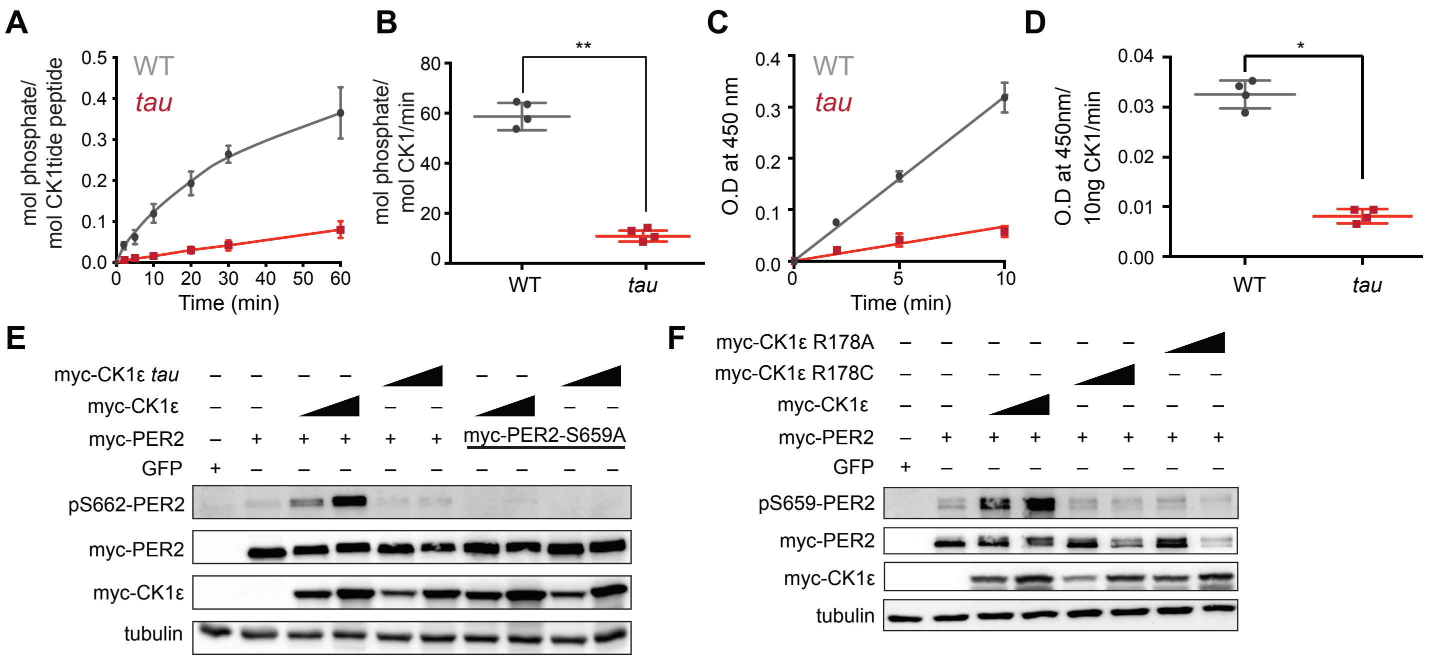


Supplemental Figure 1

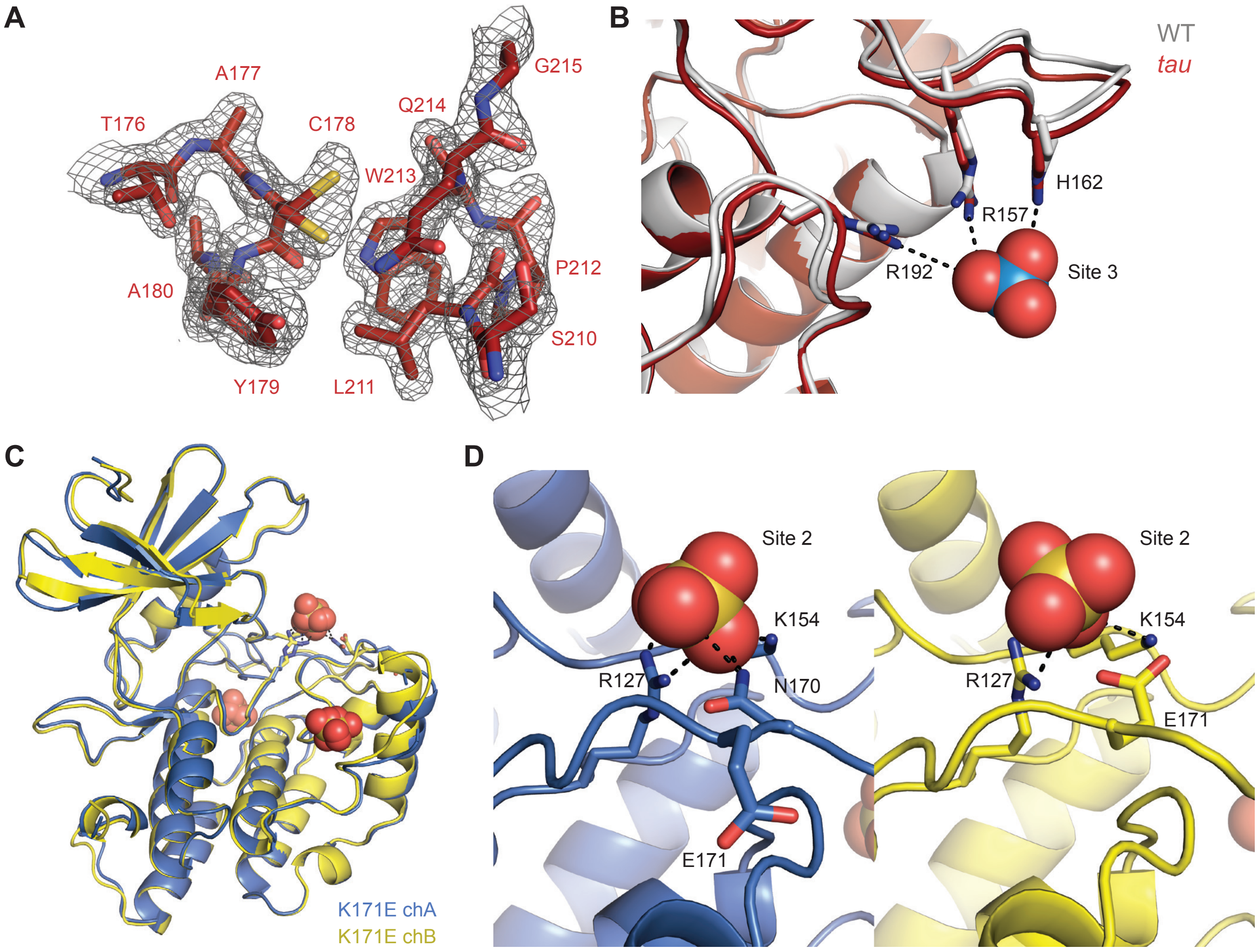


Supplemental Figure 2

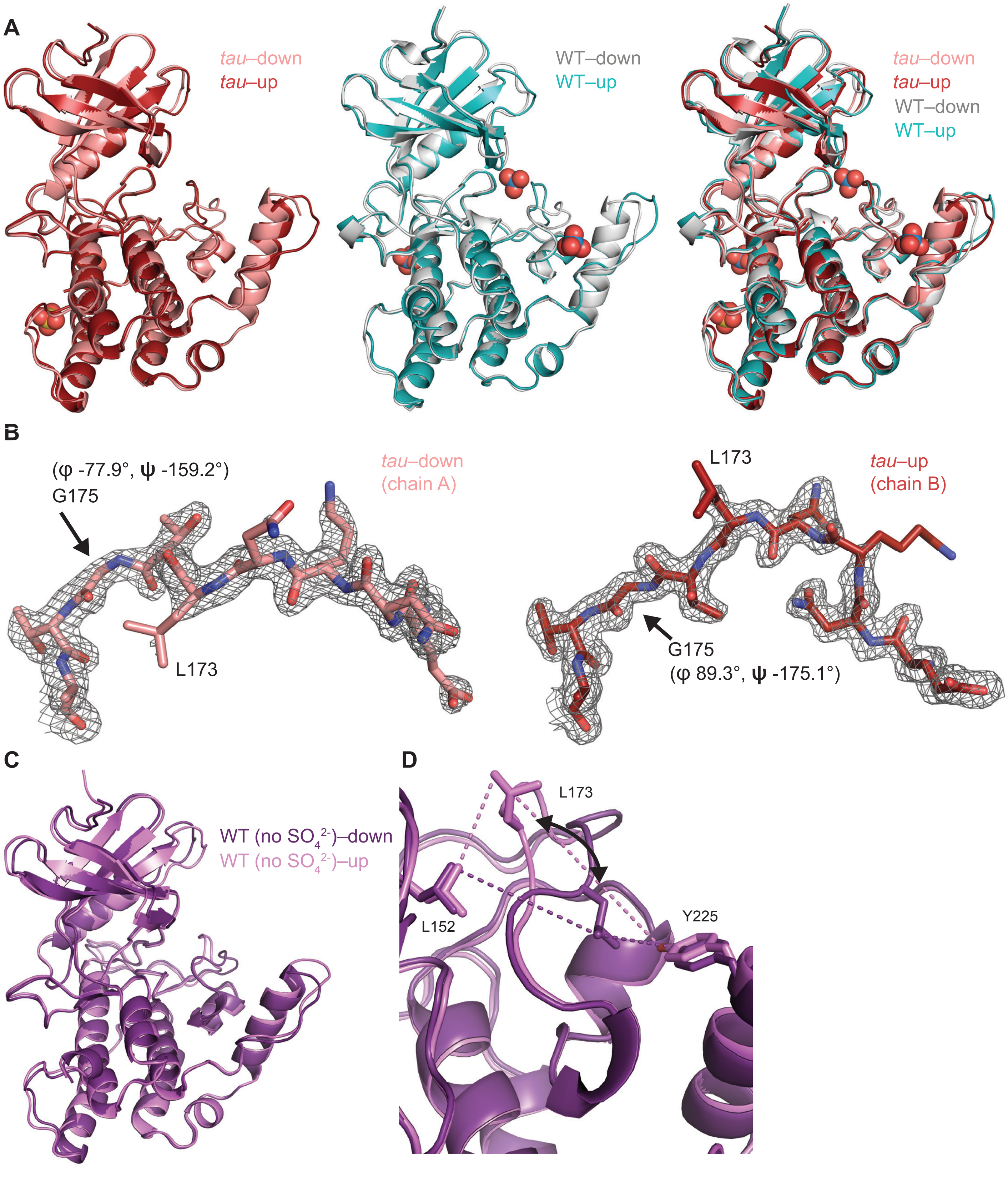


Supplemental Figure 3

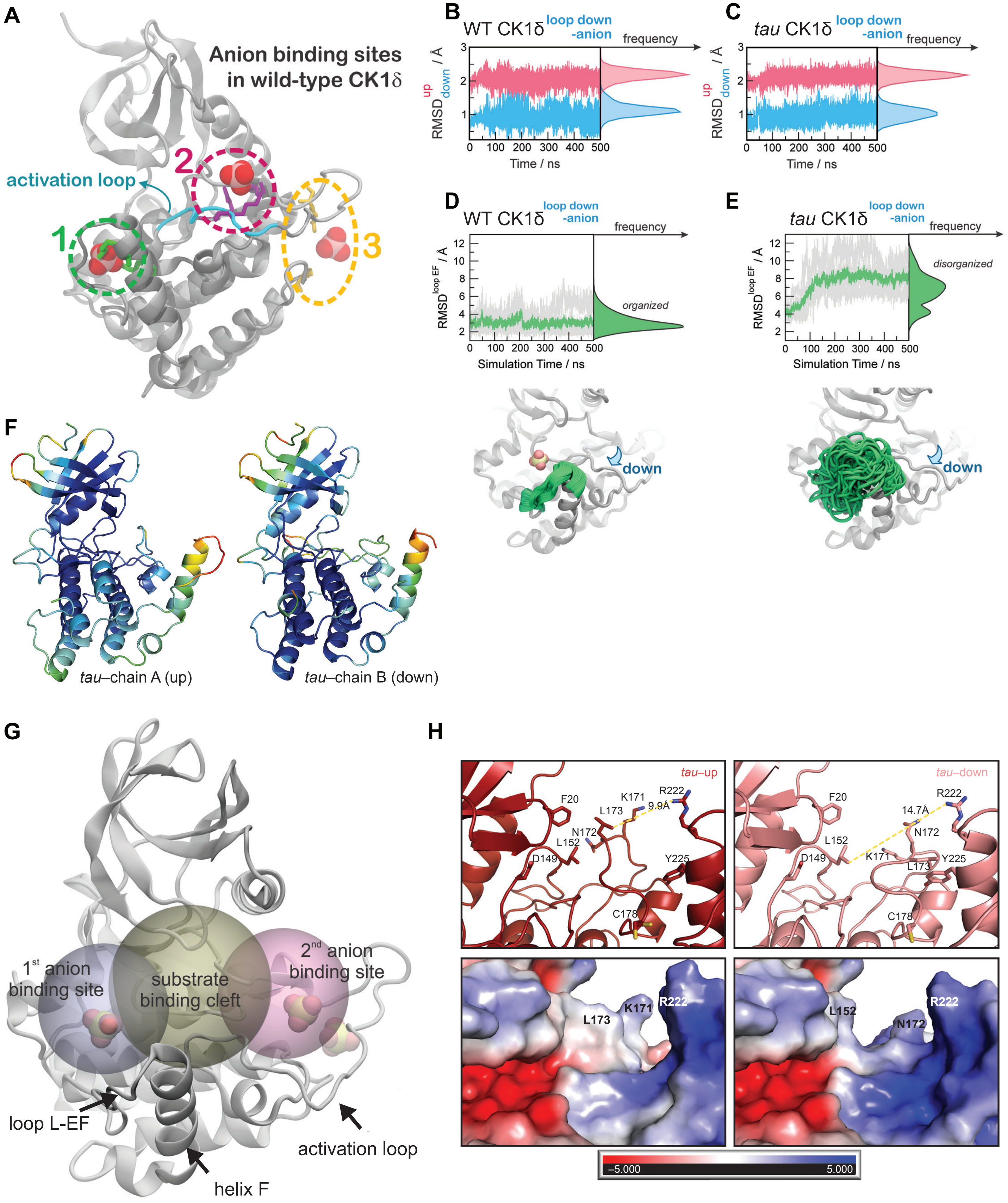


Supplemental Figure 4

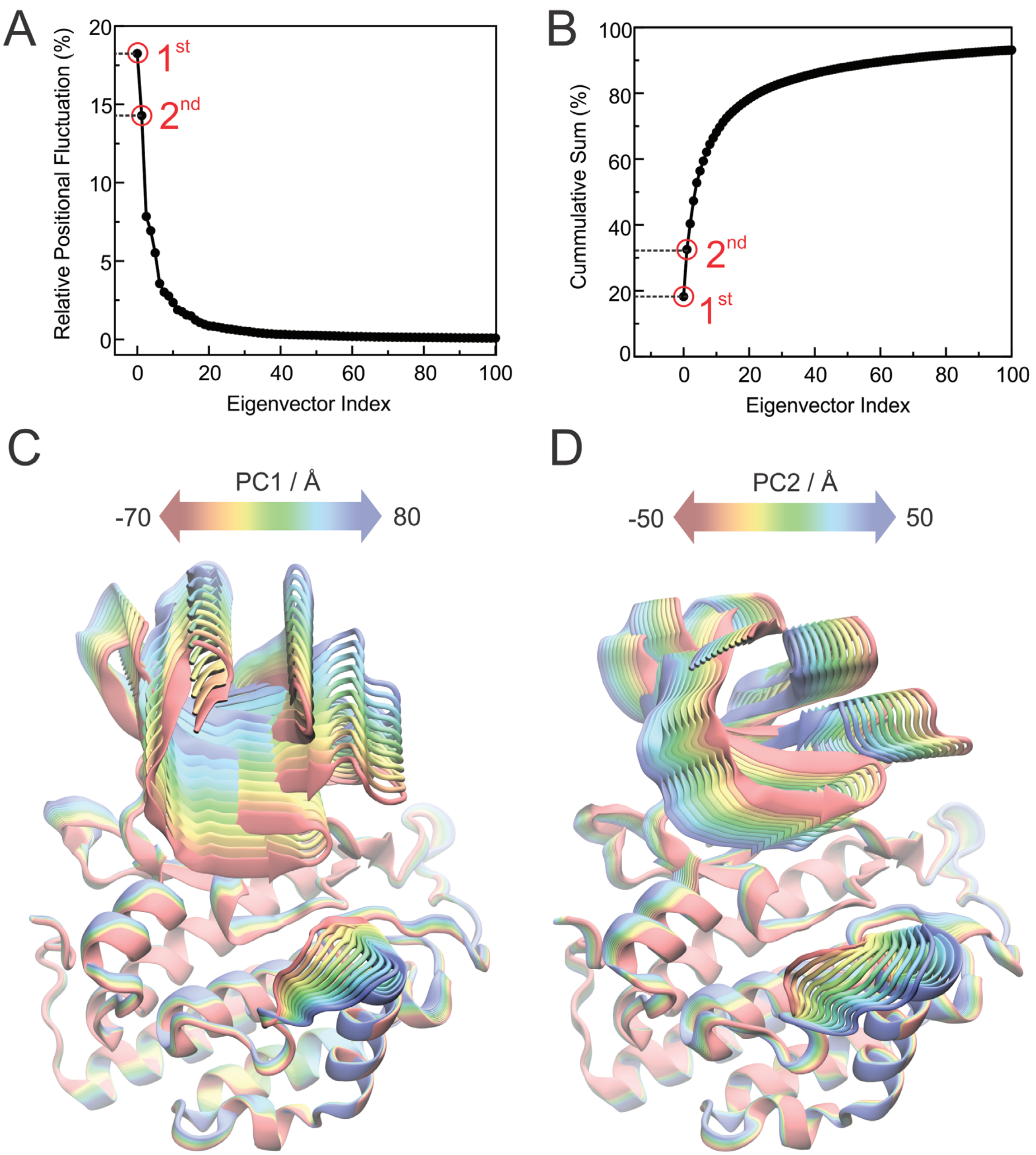


Supplemental Figure 5

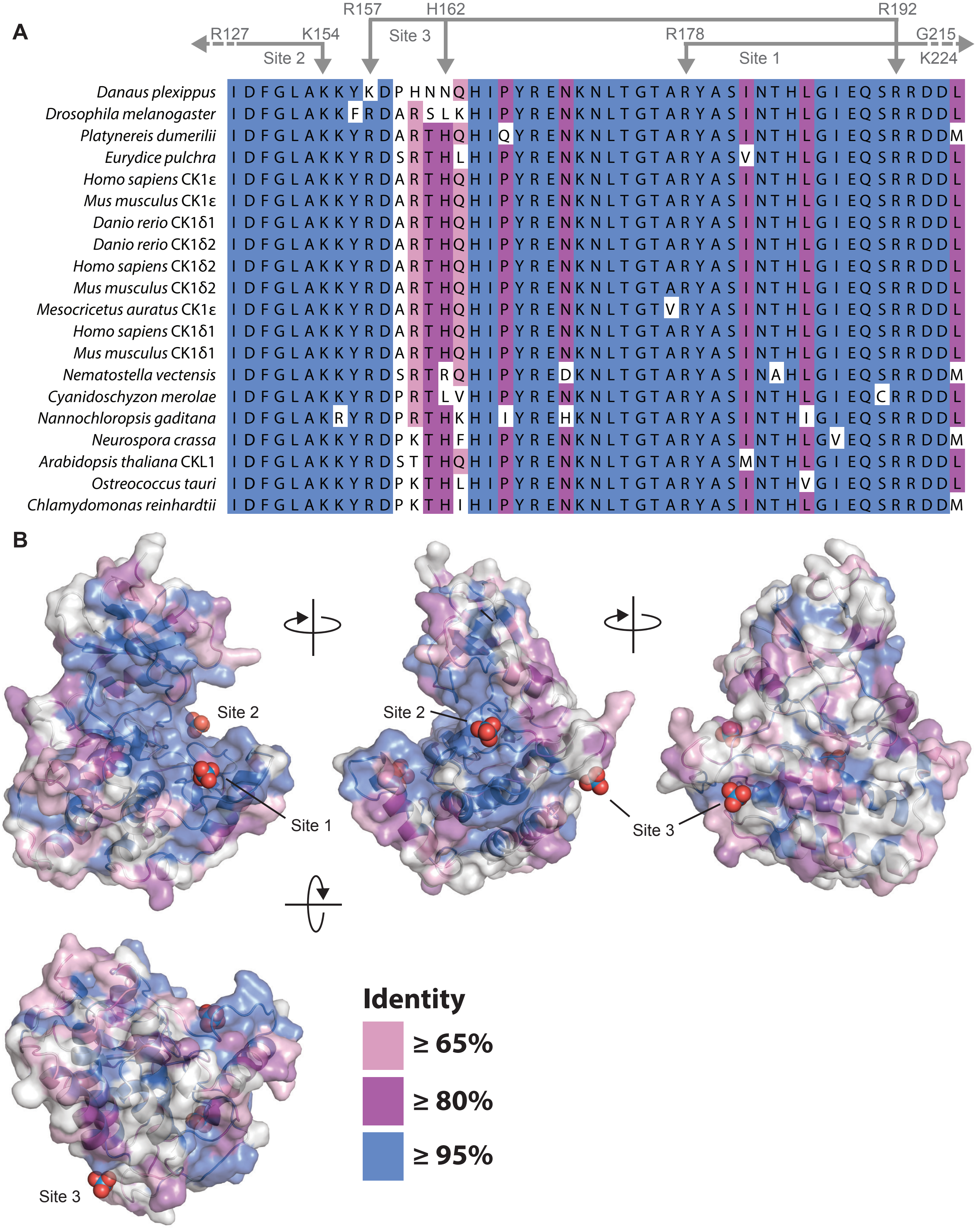


Supplemental Figure 6

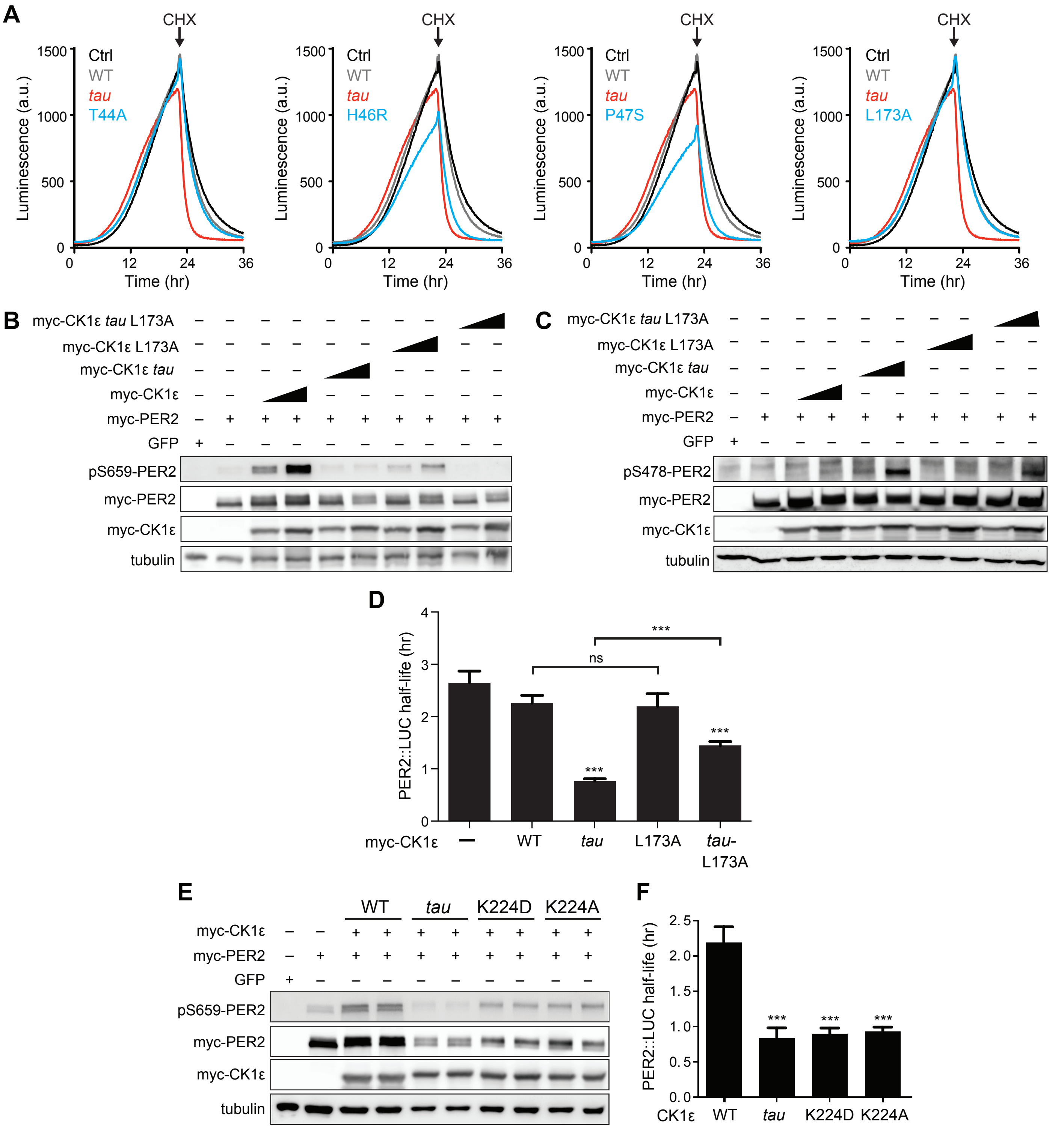


Supplemental Figure 7

**Supplemental Figure 1 (relates to Figure 1)**
A, Kinase assay using 20 nM CK1δ ΔC of WT or *tau* on 200 μM of the synthetic primed substrate, CK1tide, KRRRALpSVASLPGL (n = 4 with s.d.). B, Phosphorylation rate of kinases on CK1tide (n = 4 with s.d.). Significance assessed by Student's two-sided t-test, **, $p < 0.01$. C, ELISA-based kinase assay on 200 μM of the mouse FASP peptide detected with the anti-pS659 antibody with 10 ng of CK1δ ΔC WT or *tau* (n = 4 with s.d.). D, Rate of phosphorylation measured by optical density (O.D.) at 450 nm at the indicated timepoints (n = 4 with s.d.). Significance assessed by unpaired Student's two-sided t-test: *, $p < 0.05$. E, Western blot of sequential phosphorylation of the FASP at S662 on mouse myc-PER2 in HEK293 cell lysates after transfection with indicated expression plasmids. Representative blot from n = 3 shown. Wedge, 10 or 50 ng of myc-CK1ε plasmid used. F, Western blot of FASP priming phosphorylation at S659 on mouse myc-PER2 in HEK293 cell lysates after transfection with indicated expression plasmids as in panel E. Representative blot from n = 3 shown.

**Supplemental Figure 2 (relates to Figure 2)**
A, 2Fo-Fc omit map of the Site 1 anion binding pocket in the *tau* mutant (maroon, PDB: 6PXN, chain A) contoured at 1 σ. B, View of the Site 1 anion binding pocket with WT (gray, PDB: 1CKJ, chain B) and *tau* (maroon, PDB: 6PXN, chain A) structures overlaid. Polar interactions that coordinate the anion are depicted with dashed black lines. Note, the anions depicted at Site 1 and 2 here are only present in the WT structure. C, Overlay of K171E CK1δ ΔC (PDB: 6PXP; chain A, blue; chain B, yellow). The three conserved anion binding sites (sulfate) are depicted. D, Side-by-side overlaid view of the Site 2 anion binding pocket in the CK1δ ΔC K171E crystal structure showing polar contacts to the bound sulfate (dashed black lines) from chain A (blue, left) and chain B (yellow, right).

**Supplemental Figure 3 (relates to Figure 3)**
A, Overlay of the kinase domain of individual chains from *tau* (left, PDB: 6PXN, chains A (maroon) and B (salmon) with $SO_4^{2-}$), WT (center, PDB: 1CKJ, chains A (cyan) and B (gray) with $WO_4^{2-}$), and a comparison of all four (right). Note: Anion binding occurs at Site 1 in both chains of the WT kinase, while anion binding at Site 2 only occurs when the activation loop switch is in the down conformation of WT (chain B, gray). B, 2Fo-Fc omit maps of the activation loop in *tau* chain A and B (residues 167-177) contoured at 1 σ. C, Overlaid view of the activation loop switch in the two chains of WT CK1δ ΔC crystallized in sulfate-free conditions (PDB: 6PXO, chains A (purple) and B (violet)). D, The position of L173 CD2 relative to either L152 CD2 or Y225 OH reports on the conformation of the activation loop in the 'loop down' (purple) or 'loop up' (violet) conformation in WT CK1δ grown in sulfate-free conditions (PDB: 6PXO).

**Supplemental Figure 4 (relates to Figure 4)**
A, Defining the localization of anion binding sites 1 (green), 2 (magenta) and 3 (yellow) for MR simulations. For reference, the loop down conformation of WT CK1δ is represented as ribbons (cyan). B-C, Stability of the activation loop assessed by the Root Mean Square Deviation (RMSD) of residues 168-175 with respect to the loop down conformation ($RMSD_{down}$, blue) or to the loop up conformation ($RMSD^{up}$, magenta) as observed in the crystal structures. For both systems, the Site 2 anion was removed computationally before beginning MD simulations. Panel A, WT $CK1δ^{loop\ down}$; B, *tau* $CK1δ^{loop\ down}$; both show RMSDs superimposed from all five MD replicas. D-E, Dynamics of the EF loop obtained from Gaussian Accelerated MD simulations in WT CK1δ (panel C) and *tau* mutant (panel D) when the activation loop in the 'down' conformation and the second anion binding site has been removed computationally. The RMSD was calculated for the backbone atoms of residues 213-224 for each independent replica (gray, n = 5) and then averaged (green). The molecular representations show crystallographic structures of the enzyme (in gray) superimposed with several snapshots of the loop L-EF

extracted from the Gaussian Accelerated MD (GaMD) simulations (in green). When present, sulfate anions are represented by van der Waals spheres. F, B-factor analysis of chain A ('loop up') and B ('loop down') from CK1δ ΔC *tau* structure (PDB: 6PXN) with spectrum coloring (blue-red, increasing flexibility). G, Volumetric analysis of the Site 1 binding site (grey sphere), substrate binding cleft (golden sphere) and Site 2 anion binding site (rose sphere) during the GaMD simulations. In each system, the volumes were calculated for snapshots extracted from the GaMD trajectories every 2 ns, using POVME 3.0 (Wagner et al., 2017). Water and ions were computationally removed prior to volume calculations. H, View of the substrate-binding channel from the Site 1 anion binding pocket (near C178) looking towards the activation loop. Top row, cartoon view of protein structures: left, *tau* chain A (activation loop up, maroon); right, *tau* chain B (activation loop down, salmon). The constriction point of the substrate-binding channel was measured from L173 CD2 (left) or L152 CD2 (right) to the NH1 of R222 (dashed yellow line). Bottom row, Adaptive Poisson-Boltzmann Server (APBS) electrostatic surface representation of the same view with key residues labeled with the electrostatic potential range (± 5 kT/e) shown below (Jurrus et al., 2018).

**Supplemental Figure 5 (relates to Figure 5)**
A, Percent of atomic fluctuations contained in each of the principal modes of motion (eigenvectors) obtained from the Principal Component Analysis of the all GaMD trajectories. The 1$^{st}$ and 2$^{nd}$ modes of motions (highlighted in red) contain 18.2% and 14.3% of the total atomic fluctuations displayed by the backbone atoms, respectively. B, cumulative sum of the relative atomic fluctuations displayed in panel A, showing that, if combined, the 1$^{st}$ and 2$^{nd}$ modes of motions contain more than 30% of the total atomic fluctuations of the backbone atoms. C, 1$^{st}$ principal mode of motion corresponds to an 'open-and-close' movement of the enzyme, achieved mainly by dislocation of the N-terminal lobe (N-lobe) with respect to the top of the helix F. D, 2$^{nd}$ principal mode of motion corresponds to a twisting movement of the N-lobe with respect to the top of helix F and significant rearrangement of loop L-EF, which can either be extended or collapsed.

**Supplemental Figure 6 (relates to Figure 6)**
A, Alignment of the central catalytic DFG motif and activation loop of CK1ε and CK1δ (including isoforms δ1 and δ2 that differ only in the last 15 amino acids (Fustin et al., 2018; Narasimamurthy et al., 2018)) that have been implicated in circadian regulation of the following species: *Danaus plexippus* (Reppert et al., 2016); *Drosophila melanogaster* (Kloss et al., 1998; Price et al., 1998); *Platynereis dumerilii* (Zantke et al., 2013); *Eurydice pulchra* (Zhang et al., 2013); *Homo sapiens* (Beale et al., 2019; Toh et al., 2001; Xu et al., 2005; Xu et al., 2007); *Mus musculus* (Fustin et al., 2018; Meng et al., 2008); *Danio rerio* (Smadja Storz et al., 2013), *Mesocricetus auratus* (Ralph and Menaker, 1988); *Nematostella vectensis* (Oren et al., 2015); *Cyanidoschyzon merolae* (Matsuzaki et al., 2004); *Nannochloropsis gaditana* (Poliner et al., 2019); *Neurospora crassa* (Gorl et al., 2001); *Arabidopsis thaliana* (Uehara et al., 2019); *Ostreococcus tauri* (van Ooijen et al., 2013); and *Chlamydomonas reinhardtii* (Boesger et al., 2012, 2014). When only one CK1δ/ε-like homolog was identified in an organism, no gene name is shown in the alignment. Coloring indicates the degree of conservation, with ≥ 95% identity in blue, ≥ 80% identity in purple, ≥65% identity in pink, and <65% identity in white. Residues that coordinate anion binding on CK1 are indicated above in gray. B, Conservation from the kinase domain alignment mapped onto the WT CK1δ kinase domain (PDB: 1CKJ, chain B). The binding sites for three highly conserved anion are indicated.

**Supplemental Figure 7 (relates to Figure 7)**
A, Representative real-time luminescence data for PER2::LUC stability in HEK293 cells transfected with mouse myc-PER2::LUC plus empty vector (black) or myc-CK1ε WT (gray) or

mutants (red) as indicated (n = 4). 40 µg/mL cycloheximide (CHX) added 24 hours post-transfection (arrow). B, Western blot of FASP priming phosphorylation at S659 on mouse myc-PER2 in HEK293 cell lysates after transfection with indicated myc-CK1ε expression plasmids. Representative blot from n = 3 shown. C, Western blot of Degron phosphorylation at S478 on mouse myc-PER2 in HEK293 cell lysates after transfection with indicated myc-CK1ε expression plasmids. Representative blot from n = 3 shown. D, Quantification of PER2::LUC half-life from transfection assays (as in panel A) with different myc-CK1ε mutants. Data represent mean ± s.d. (n = 4). E, Western blot of FASP priming phosphorylation at S659 on mouse myc-PER2 in HEK293 cell lysates after transfection with indicated myc-CK1ε expression plasmids. Representative blot from n = 3 shown. F, Quantification of PER2::LUC half-life from transfection assays (as in panel A) with myc-CK1ε mutants from panel E. Data represent mean ± s.d. (n = 4).

**Supplemental Movie 1 (relates to Figure 5)**
**Principal Component Analysis of CK1δ normal modes**
1$^{st}$ principal mode of motion corresponds to an ‘open-and-close’ movement of the enzyme, achieved mainly by dislocation of the N-terminal lobe (N-lobe) with respect to the top of the helix F. 2$^{nd}$ principal mode of motion corresponds to a twisting movement of the N-lobe with respect to the top of helix F and significant rearrangement of loop L-EF, which can either be extended or collapsed.

**Supplemental Table 1 – X-ray crystallography data collection and refinement statistics**

| | CK1δ ΔC *tau* R178C | CK1δ ΔC No Sulfate | CK1δ ΔC K171E |
|---|---|---|---|
| **PDB** | 6PXN | 6PXO | 6PXP |
| ***Data collection*** | | | |
| Space group | $P2_1$ | $P2_1$ | $P2_1$ |
| Resolution | 64.82-1.55 (1.58-1.55) | 65.24-2.00 (2.11-2.00) | 47.38 – 2.35 (2.43-2.35) |
| a, b, c | 50.08, 129.64, 51.86 | 49.91, 130.49, 51.16 | 50.06, 130.35, 51.65 |
| $\alpha, \beta, \gamma$ | 90, 113.41, 90 | 90, 112.75, 90 | 90, 113.47, 90 |
| $R_{merge}$ | 6.3 (60.1) | 10.5 (104.2) | 12.8 (65.2) |
| $R_{pim}$ | 2.8 (27.0) | 4.5 (46.8) | 7.7 (40.4) |
| Total reflections | 588739 (27527) | 252539 (34017) | 109702 (10773) |
| Unique reflections | 85497 (4108) | 40494 (5884) | 25271 (2470) |
| $I/\sigma$ | 13.1 (2.5) | 11.2 (1.7) | 6.7 (2.1) |
| CC1/2 | 0.99(0.89) | 0.99 (0.61) | 0.99 (0.77) |
| Completeness | 97.8(95.3) | 99.6 (99.7) | 99.9 (99.9) |
| Redundancy | 6.9 (6.7) | 6.2 (5.8) | 4.3 (4.4) |
| Wilson B-Factor | 19 | 28 | 32 |
| ***Refinement*** | | | |
| Resolution | 44.68-1.55 | 47.18-2.0 | 42.5-2.35 |
| $R_{work}/R_{free}$ | 17.9/20.6 | 17.9/23.7 | 20.0/24.3 |
| No. of Atoms | 5057 | 5012 | 4922 |
| Protein | 4735 | 4727 | 4757 |
| Water | 297 | 285 | 115 |
| Ligands | 25 | - | 50 |
| ***RMS deviation*** | | | |
| Bond lengths | 0.01 | 0.008 | 0.009 |
| Bond angles | 1.2 | 0.94 | 1.15 |
| Ramachandran Favored/Outliers | 100.0/0.0 | 100.0/0.0 | 100/0.0 |
| Average B-Factor | 34 | 41 | 41 |

Values in parentheses are for highest resolution shell

**Supplemental Table 2 Enzymatic efficiency of CK1δ ΔC (WT and mutants)**

| CK1δ ΔC | FASP $k_{cat}/K_m$ ($M^{-1}$ $s^{-1}$) | Degron $k_{cat}/K_m$ ($M^{-1}$ $s^{-1}$) | FASP/Degron Efficiency Ratio |
|---|---|---|---|
| wild-type | 0.0576 ± 0.0012 | 0.0322 ± 0.0009 | 1.77 ± 0.09 |
| T44A | 0.0447 ± 0.0007 | 0.0332 ± 0.0017 | 1.37 ± 0.08 |
| H46R | 0.0237 ± 0.0005 | 0.0235 ± 0.0009 | 1.01 ± 0.08 |
| P47S | 0.0349 ± 0.0008 | 0.0312 ± 0.0020 | 1.14 ± 0.09 |
| R127E | 0.0464 ± 0.0016 | 0.0479 ± 0.0011 | 0.969 ± 0.05 |
| K154E | 0.0381 ± 0.0008 | 0.0350 ± 0.0021 | 1.09 ± 0.05 |
| K171E | 0.0425 ± 0.0008 | 0.0363 ± 0.0005 | 1.17 ± 0.03 |
| L173A | 0.0358 ± 0.0037 | 0.0276 ± 0.0012 | 1.30 ± 0.06 |
| *tau* (R178C) | 0.0350 ± 0.0013 | 0.0368 ± 0.0008 | 0.918 ± 0.02 |
| *tau* L173A | 0.0203 ± 0.0007 | 0.0198 ± 0.0002 | 1.02 ± 0.03 |
| K224D | 0.0345 ± 0.0010 | 0.0319 ± 0.0009 | 1.08 ± 0.01 |

$k_{cat}/K_m$ data are represented as mean ± s.d. from n = 3 assays.

**Supplemental Table 3 – Survey of anion binding and activation loop conformation in CK1 family member structures**

| PDB code | Species | Isoform | Res. (Å) | Space group | Site 1 Occ.† | Site 2 Occ.† | A-loop switch conformation | Mutation or ligand |
|---|---|---|---|---|---|---|---|---|
| 1CKI | Rat | CK1δ | 2.30 | $P\ 2_1\ 2_1\ 2_1$ | no | no | down | |
| 1CKJ | Rat | CK1δ | 2.46 | $P\ 2_1\ 2_1\ 2_1$ | yes | yes/no | down/up | |
| 4HGT | Human | CK1δ | 1.80 | $P\ 1\ 2_1\ 1$ | no | no | down | ligand |
| 4HNF | Human | CK1δ | 2.07 | C 1 2 1 | no | no | down | ligand |
| 3UYS | Human | CK1δ | 2.30 | P1 | yes | yes/no | down | |
| 3UYT | Human | CK1δ | 2.00 | P1 | yes | yes/no | down/poor density | ligand |
| 3UZP | Human | CK1δ | 1.94 | $P\ 1\ 2_1\ 1$ | no | no | down | ligand |
| 4TWC | Human | CK1δ | 1.70 | $P\ 1\ 2_1\ 1$ | yes | yes | down | ligand |
| 4JJR | Mouse | CK1δ | 2.41 | $P\ 1\ 2_1\ 1$ | yes | yes | down | |
| 4KB8 | Human | CK1δ | 1.95 | P1 | yes | yes/no | down/poor density | ligand |
| 4KBA | Human | CK1δ | 1.98 | P1 | yes | yes/no | down/poor density | ligand |
| 4KBC | Human | CK1δ | 1.98 | P1 | yes | yes | down | ligand |
| 4KBK | Human | CK1δ | 2.10 | P1 | yes | yes | down | ligand |
| 4TW9 | Human | CK1δ | 2.40 | $P\ 1\ 2_1\ 1$ | yes | yes | down | ligand |
| 4TN6 | Human | CK1δ | 2.41 | $P\ 1\ 2_1\ 1$ | yes | yes | down | ligand |
| 5IH4 | Human | CK1δ | 1.90 | $P\ 3_1\ 2\ 1$ | yes | yes | down | |
| 5IH5 | Human | CK1δ | 2.25 | $P\ 3_1\ 2\ 1$ | yes | yes | down | ligand |
| 5IH6 | Human | CK1δ | 2.30 | $P\ 3_1\ 2\ 1$ | yes | yes | down | ligand |
| 5W4W | Human | CK1δ | 1.99 | P1 | yes | yes/no | down | ligand |
| 5MQV | Human | CK1δ | 2.15 | C 1 2 1 | yes | yes/no | down | ligand |
| 5X17 | Human | CK1δ | 2.00 | $P\ 1\ 2_1\ 1$ | yes | yes | down | ADP |
| 5OKT | Human | CK1δ | 2.13 | $P\ 1\ 2_1\ 1$ | yes | yes | down | ligand |
| 6GZM | Human | CK1δ | 1.59 | $P\ 1\ 2_1\ 1$ | yes | yes | down | ligand |
| 6F1W | Human | CK1δ | 1.86 | $P\ 1\ 2_1\ 1$ | yes | yes | down | ligand |
| 6F26 | Human | CK1δ | 1.83 | $P\ 1\ 2_1\ 1$ | yes | yes | down | ligand |
| 6PXN | Human | CK1δ | 1.55 | $P2_1$ | no | yes/no | down/up | R178C |
| 6PXO | Human | CK1δ | 2.00 | $P2_1$ | no | no | down/up | WT |
| 6PXP | Human | CK1δ | 2.35 | $P2_1$ | yes | yes | down | K171E |
| 4HOK | Human | CK1ε | 2.77 | C 1 2 1 | yes | no | down | |
| 5X18 | *S. cerevisiae* | CK1 | 1.80 | $P\ 1_2\ 1\ 1$ | no | no | down | |
| 1CSN | *S.pombe* | CK1 | 2.00 | $P\ 3_2\ 2\ 1$ | yes | yes | down | Mg-ATP |
| 2CSN | *S.pombe* | CK1 | 2.50 | $P\ 3_2\ 2\ 1$ | yes | yes | down | ligand |
| 1EH4 | *S.pombe* | CK1 | 2.80 | $P\ 6_1$ | yes | yes | down | ligand |
| 4XH0 | *C.glabrata* | Hrr25 | 1.99 | $P\ 2_1\ 2\ 2_1$ | yes | yes | down | ADP |
| 4XHG | *C.glabrata* | Hrr25 | 2.15 | $P\ 2_1\ 2\ 2_1$ | no | no | down | ADP |
| 4XHH | *C.glabrata* | Hrr25 | 2.91 | $P\ 2_1\ 2\ 2_1$ | yes | yes | down | |
| 4XHL | *C.glabrata* | Hrr25 | 3.1 | $P\ 2_1\ 2\ 2_1$ | yes | yes | down | K38R |
| 2CMW | Human | CK1ɣ1 | 1.75 | $P\ 2_1\ 2_1\ 2_1$ | yes | no | down | |
| 2C47 | Human | CK1ɣ2 | 2.40 | $P\ 2_1\ 2_1\ 2_1$ | no | no | down | ligand |
| 2CHL* | Human | CK1ɣ3 | 1.95 | $P\ 3_1\ 2\ 1$ | yes | yes | down | ligand |
| 3SV0 | *O. sativa* | CK1-like | 2.00 | C 1 2 1 | no | no | down | |

†Reflects anion occupancy (Occ.) at Site 1 or Site 2

*Other CK1ɣ3 structures with ligands (all act. loop switch down): 2IZR, 2IZS, 2IZT, 2IZU, 4G16, 4HGL, 4HGS

**Supplemental Table 4 – CK1 family alleles and their circadian phenotypes**

| Organism | Gene | Allele | Residue(s) | Studied in | Circadian Phenotype |
|---|---|---|---|---|---|
| Syrian Hamster | CK1ε | *tau* | R178C | Syrian Hamster | Short period (Ralph and Menaker, 1988) |
| Mouse | CK1ε | *tau* | R178C | Mouse | Short period (Lowrey et al., 2000) |
| *Drosophila* | DBT | *dbt^tau^* | R178C | *Drosophila* | Short period (Fan et al., 2009) |
| Human | CK1ε | | S408N | n.d. | Protective against DSPS (Takano et al., 2004) |
| Human | CK1δ | | T44A | Mouse | Short period (Xu et al., 2005) |
| | CK1δ | | T44A | *Drosophila* | Long period (Xu et al., 2005) |
| Human | CK1δ | | H46R | *In vitro* | Low activity (Brennan et al., 2013) |
| Mouse | CK1δ | | K224D | Mouse | Short period (Shinohara et al., 2017) |
| *Drosophila* | DBT | $dco^{2}$ | G175S | *Drosophila* | n.d. (Zilian et al., 1999) |
| *Drosophila* | DBT | $dco^{3}$ | R4C, E47K | *Drosophila* | n.d. (Zilian et al., 1999) |
| *Drosophila* | DBT | $dco^{18}$ | S181F | *Drosophila* | n.d. (Zilian et al., 1999) |
| *Drosophila* | DBT | $dbt^{S}$ | P47S | *Drosophila* | Short period (Kloss et al., 1998; Price et al., 1998) |
| *Drosophila* | DBT | $dbt^{L}$ | M80I | *Drosophila* | Long period (Kloss et al., 1998; Price et al., 1998) |
| *Drosophila* | DBT | $dbt^{G}$ | R127H | *Drosophila* | Long period (Suri et al., 2000) |
| *Drosophila* | DBT | $dbt^{H}$ | T44I | *Drosophila* | Long period (Suri et al., 2000) |
| *Drosophila* | DBT | $dbt^{AR}$ | H126Y | *Drosophila* | Long period (Rothenfluh et al., 2000) |
| *Drosophila* | DBT | $dbt^{K/R}$ | K38R | *Drosophila* | Long period (Muskus et al., 2007) |
| *Drosophila* | DBT | $dbt^{P}$ | 5' UTR | *Drosophila* | Arrhythmic in larva ( PER stabilized); lethal in pupae (Kloss et al., 1998; Price et al., 1998) |
| *Drosophila* | DBT | $dbt^{EY02910}$ | 5' UTR | *Drosophila* | Arrhythmic or long period (Zheng et al., 2014) |

**Abbreviations**: DSPS, Delayed Sleep Phase Disorder; n.d., not determined

**Supplemental Table 5 – Simulated systems**

| System | Activation loop conformation | Presence of $SO_4^{2-}$ ions | | | Chain | PDB |
|---|---|---|---|---|---|---|
| | | Site 1 | Site 2 | Site 3 | | |
| WT CK1$\delta^{loop\ down}$ | Down | Yes | Yes | Yes | B | 1CKJ |
| WT CK1$\delta^{loop\ up}$ | Up | Yes | No | Yes | A | 1CKJ |
| *tau* CK1$\delta^{loop\ down}$ | Down | No | Yes | Yes | B | 6PXN |
| *tau* CK1$\delta^{loop\ up}$ | Up | No | No | Yes | A | 6PXN |
| WT CK1$\delta^{loop\ down}{}_{-anion}$ | Down | Yes | No* | Yes | B | 1CKJ |
| *tau* CK1$\delta^{loop\ down}{}_{-anion}$ | Down | No | No* | Yes | B | 6PXN |

* $SO_4^{2-}$ ions were computationally removed

The nomenclature of the anion binding sites is defined in Figure S4.

### Materials and Methods

#### Cell culture, reagents and transfection

myc-mPer2, myc-mPer2 S659A, mPer2::Luc, and myc-CK1ε expression plasmids were described previously (Eide et al., 2005; Eng et al., 2017; Zhou et al., 2015). Mutations of the kinase domain were introduced by Quikchange site-directed mutagenesis (Stratagene) and validated by sequencing.

HEK293 cells (from American Type Culture Collection) were cultured in Dulbecco's Modified Eagle's Medium (DMEM, Gibco) supplemented with 10% FBS (Gibco), 50 units/mL penicillin, 50 µg/mL streptomycin (Invitrogen) and maintained at 37°C in a 5% $CO_2$ environment. Cells were transfected using Lipofectamine 2000 transfection reagent (Life Technologies) following the manufacturer's instructions. For transfections titrating expression of myc-CK1ε, either 10 or 50 ng of plasmid was used; total plasmid DNA of either 1 or 2 µg was used for each well of a 12 or 6-well-plate respectively.10 µM MG132 was added to cultures 24 hours prior to harvest to prevent proteasomal degradation for experiments shown in Figure 7A and B.

#### PER2::LUC half-life measurement

Mouse PER2::LUC expression plasmids (10 ng) were transiently transfected alone or with myc-CK1ε (100 ng) in 35 mm dishes of HEK293 cells in phenol red-free DMEM in the presence of 100 mM D-luciferin (122799, PerkinElmer), 10 mM HEPES and 1.2 g/L sodium bicarbonate. Dishes were sealed with 40 mm cover glasses and vacuum grease, and incubated in the LumiCycle (Actimetrics). The next day, 40 µg/mL cycloheximide (Sigma) was added per 35 mm dish. Luminescence data were used to calculate PER2::LUC half-life in Prism (GraphPad) using one-phase decay algorithm as described previously (Zhou et al., 2015). Briefly, half-lives were calculated using the one-phase decay algorithm in Prism (GraphPad) using the raw luciferase activity, beginning from the point of cycloheximide addition to the plateau at minimum luciferase activity (n = 4).

#### SDS-PAGE and western blotting

Whole cell extracts of transfected HEK293 cells lysed on ice with cell lysis buffer (50 mM Tris-HCl pH 8.0, 150 mM NaCl, 1% (vol/vol) Nonidet P-40 and 0.5% deoxycholic acid containing Complete Protease Inhibitors (Roche) and PhosStop Phosphatase Inhibitors (Roche)) were analyzed by denaturing SDS-PAGE gel, which was transferred on PVDF membrane (Immobilon, Millipore). The blot was probed using the indicated primary antibodies: anti-myc (9E10) (sc-40, Santa Cruz Biotechnology) and anti-tubulin (ab52623, Abcam) were purchased from commercial providers, while rabbit polyclonal antibodies were generated against phospho-Ser478, phospho-Ser659 or phospho-Ser662 of mouse PER2 and purified against the phosphopeptides by Abfrontier (Young In Frontier Co.). The phosphopeptides for phospho-Ser478 and phospho-Ser659 have been described elsewhere (Narasimamurthy et al., 2018; Zhou et al., 2015); the phosphopeptide KAESVVpSLTSQ-Cys was used to generate the phospho-Ser662 antibody. HRP-conjugated goat secondary antibodies for anti-rabbit (1706515, Bio-Rad) and anti-mouse (1706516, Bio-Rad) were with standard ECL reagents (Thermo Fisher Scientific). Densitometric analysis of western blot bands was performed using ImageJ software (National Institutes of Health).

**Expression and purification of recombinant proteins**

All proteins were expressed from a pET22-based vector in *Escherichia coli* Rosetta2 (DE3) cells based on the Parallel vector series (Sheffield et al., 1999). The extended wild-type FASP peptide (residues 645-687) and Degron peptide (residues 475-505) from human PER2, and a short wild-type mouse FASP peptide (residues 642-666) all contain an N-terminal WRKKK polybasic motif for *in vitro* kinase assays and a tryptophan for UV detection during purification. All peptides were expressed downstream of an N-terminal TEV-cleavable His-NusA tag. Human CK1δ catalytic domains (CK1δ ΔC, residues 1-317) were all expressed in Rosetta2 (DE3) cells with a TEV-cleavable His-GST tag. Mutations were made using standard site-directed mutagenesis protocols and validated by sequencing. All proteins and peptides expressed from Parallel vectors have an additional N-terminal vector artifact of "GAMDPEF" remaining after TEV cleavage. Cells were grown in LB media (for natural abundance growths) or M9 minimal medium with the appropriate stable isotopes ($^{15}N$, $^{13}C$ for NMR, as done before (Narasimamurthy et al., 2018)) at 37 °C until the $O.D._{600}$ reached ~0.8; expression was induced with 0.5 mM IPTG, and cultures were grown for approximately 16-20 hours more at 18 °C.

For CK1δ kinase domain protein preps, cells were lysed in 50 mM Tris pH 7.5, 300 mM NaCl, 1 mM TCEP, and 5% glycerol using a high-pressure extruder (Avestin). HisGST-CK1δ ΔC fusion proteins were purified using Glutathione Sepharose 4B resin (GE Healthcare) using standard approaches and eluted from the resin using Phosphate Buffered Saline with 25 mM reduced glutathione. $His_6$-TEV protease was added to cleave the His-GST tag from CK1δ ΔC at 4 °C overnight. Cleaved CK1δ ΔC was further purified away from His-GST and His-TEV using Ni-NTA resin (Qiagen) and subsequent size exclusion chromatography on a HiLoad 16/600 Superdex 75 prep grade column (GE Healthcare) in 50 mM Tris pH 7.5, 200 mM NaCl, 5 mM BME, 1 mM EDTA, and 0.05% (vol/vol) Tween 20. Purified CK1δ ΔC proteins used for *in vitro* kinase assays were buffer exchanged into storage buffer (50 mM Tris pH 7.5, 100 mM NaCl, 1 mM TCEP, 1 mM EDTA, and 10% glycerol) using an Amicron Ultra centrifugal filter (Millipore) and frozen as small aliquots in liquid nitrogen for storage at -80 °C.

For PER2 peptide preps, cells were lysed in 50 mM Tris pH 7.5, 500 mM NaCl, 2 mM TCEP, 5 % glycerol and 25 mM imidazole using a high-pressure extruder (Avestin). His-NusA-FASP or His-NusA -Degron fusion proteins were purified using Ni-NTA resin using standard approaches and eluted from the resin using 50 mM Tris pH 7.5, 500 mM NaCl, 2 mM TCEP, 5 % glycerol and 250 mM imidazole. His-TEV protease was added to cleave the $His_6$-NusA tag from the PER2 peptides at 4 °C overnight. The cleavage reaction was subsequently concentrated and desalted into low imidazole lysis buffer using a HiPrep 26/10 Desalting column. Peptides were purified away from His-NusA and His-TEV using Ni-NTA resin with 50 mM Tris pH 7.5, 500 mM NaCl, 2 mM TCEP, 5 % glycerol and 25 mM imidazole. Peptides were purified by size exclusion chromatography on a HiLoad 16/600 Superdex 75 prep grade column, using NMR buffer (25 mM MES pH 6.0, 50 mM NaCl, 2 mM TCEP, 1 mM EDTA, 11 mM $MgCl_2$) or 1x kinase buffer (25 mM Tris pH 7.5, 100 mM NaCl, 10 mM $MgCl_2$, and 2 mM TCEP) for NMR or ADP-Glo kinase assays, respectively.

**Radioactive and ELISA-based kinase assays**

Mouse PER2 FASP region peptides (primed, RKKKTEVSAHLSSLTLPGKAEpSVVSLTSQ, or unprimed, RKKKTEVSAHLSSLTLPGKAESVVSLTSQ), mouse PER2 Degron peptide (RKKKPHSGSSGYGSLGSNGSHEHMSQTSSSDSN, from (Isojima et al., 2009)) and the CK1tide peptide (KRRRALpSVASLPGL, from (Isojima et al., 2009; Shinohara et al., 2017)) were synthesized and purified to 95% or higher (SABio).

For the radioactive kinase assay, two independent reaction mixtures of 50 µL containing 200 µM of the FASP or Degron peptides in reaction buffer (25 mM Tris pH 7.5, 7.5 mM $MgCl_2$, 1 mM DTT, 0.1 mg/mL BSA) were preincubated for 5 minutes with or without 20 nM CK1δ ΔC (for primed FASP substrate) or 200 nM CK1δ ΔC (for unprimed FASP or Degron) and the reaction was started by addition of 750 µM of UltraPure ATP (Promega) containing 1-2 µCi of ɣ-$^{32}$p ATP (Perkin Elmer). After incubation of the reaction mix at 30 °C, an 8 µL aliquot of the reaction mix was transferred to P81 phosphocellulose paper (Reaction Biology Corp) at the indicated timepoints. The P81 paper was washed three times with 75 mM of orthophosphoric acid and once with acetone. The air-dried P81 paper was counted for $P_i$ incorporation using a scintillation counter (Perkin Elmer) by Cherenkov counting. Results shown are from four independent assays.

For the ELISA kinase assay, unprimed FASP peptide was diluted to 2 µg/mL in Carbonate buffer, pH 9.5 (0.1 M sodium carbonate) and coated onto a 96 well plate (100 µL/well). The next day, wells were washed three times with wash buffer (PBS with 0.05% Tween, PBS-T) and once with kinase buffer (25 mM Tris pH 7.5, 5 mM beta glycerol phosphate, 2 mM DTT and 0.1 mM sodium orthovanadate). Reaction mixture (50 µL) containing 10 ng of CK1δ ΔC purified protein in the kinase buffer including 10 mM $MgCl_2$ and 200 µM ATP was added onto each well and the plate was incubated at 30 °C for 1 hour. Next, the reaction mixture was removed and the wells were washed with three times with wash buffer and incubated with blocking buffer (PBS-T with 5% BSA) for 1 hour at room temperature. Subsequently, wells were incubated with pS659 Ab, anti-rabbit antibody conjugated to Biotin and Streptavidin-HRP for 1 hour at room temperature with a washing step after each incubation as above. For signal detection, TMB (1-Step Ultra TMB-ELISA, Thermo Scientific) was added, incubated for color development at room temperature and stopped with the addition of STOP solution (Thermo Scientific). The plate was read at 450 nm using an xMark Spectrophotometer plate reader (Biorad). Results shown are from four independent assays.

**NMR-based kinase assay**

NMR spectra were collected on a Varian INOVA 600 MHz or a Bruker 800 MHz spectrometer equipped with a $^{1}$H, $^{13}$C, $^{15}$N triple resonance z-axis pulsed-field-gradient cryoprobe. Spectra were processed using NMRPipe (Delaglio et al., 1995) and analyzed using CCPNmr Analysis (Vranken et al., 2005). Backbone assignments were obtained previously for the mouse FASP (BMRB entry: 27306) (Narasimamurthy et al., 2018). NMR kinase reactions were performed at 30 ˚C with 0.2 mM $^{15}$N-mouse FASP, 2.5 mM ATP and 1 µM CK1d ΔC (WT or *tau*). SOFAST HMQC spectra (total data acquisition = 6 min) were collected at the indicated intervals for 3 hours and relative peak volumes were calculated and normalized as described previously (Narasimamurthy et al., 2018). Data analysis was performed using Prism (GraphPad), with data fit to either a one-phase exponential or linear regression.

**ADP-Glo kinase assay**

Kinase reactions were performed on the indicated peptides (Degron or extended FASP) using the ADP-Glo kinase assay kit (Promega) according to manufacturer's instructions. All reactions were performed in 30 µL volumes using 1x kinase buffer (25 mM Tris pH 7.5, 100 mM NaCl, 10 mM $MgCl_2$, and 2 mM TCEP) supplemented with ATP and PER2 substrate peptides as indicated. To determine the apparent 2$^{nd}$-order rate constants, triplicate reactions containing 10 µM substrate, 100 µM ATP, and 0.2 µM CK1δ ΔC kinase were incubated in 1x kinase buffer at room temperature for 3 hours (and repeated for n = 3 independent assays). Linearity of the reaction rate with respect to time was determined by performing larger reactions (50 µL) with wild-type and *tau* CK1δ ΔC and either the FASP or Degron substrate; 5 µL aliquots were taken and quenched with ADP-Glo reagent at discrete time points up to 3 hours (data not shown). Luminescence measurements were taken at room temperature with a SYNERGY2 microplate reader in 384-well microplates. Data analysis was performed using Excel (Microsoft) or Prism (GraphPad).

**Statistical analyses**

All statistical analyses were done using Prism (GraphPad). p-values were calculated using unpaired two-tailed Students t-tests. In all figures, * indicates $p<0.05$, ** $p<0.01$, *** $p<0.001$, **** $p<0.0001$; ns, not significant.

**Crystallization and structure determination**

Crystallization was performed by hanging-drop vapor-diffusion method at 22 °C by mixing an equal volume of CK1δ ΔC with reservoir solution. The reservoir solution for CK1δ ΔC *tau* (R178C) (8.5 mg/mL) was 50 mM sodium acetate pH 6.0, 300 mM ammonium sulfate, and 17.5% (vol/vol) PEG 2000. The reservoir solution for crystallizing CK1δ ΔC wild-type (8 mg/mL) without sulfate was 150 mM succinic acid pH 5.5 and 17% (vol/vol) PEG 3350. The reservoir solution for CK1δ ΔC K171E (7.7 mg/mL) was 50 mM sodium acetate pH 6.0, 350 mM ammonium sulfate, and 17.5% (vol/vol) PEG 3500. The crystals were looped and briefly soaked in a drop of reservoir solution and then flash-cooled in liquid nitrogen for X-ray diffraction data collection. For CK1δ ΔC *tau*, a cryopreservant of reservoir solution with 20% (vol/vol) glycerol was used. Data sets were collected at the APS beamline 23-ID-D, and the ALS beamline 8.3.1. Data were indexed, integrated and merged using the CCP4 software suite (Winn et al., 2011). Structures were determined by molecular replacement with Phaser MR (McCoy et al., 2007) using the ADP-bound structure of wild-type CK1δ ΔC (PDB: 5X17). Model building was performed with Coot (Emsley et al., 2010) and structure refinement was performed with PHENIX (Adams et al., 2011). All structural models and alignments were generated using PyMOL Molecular Graphics System 2.0 (Schrödinger).

**Molecular dynamics**

*Initial structures:* To simulate *tau* CK1δ, we used the crystallographic structure reported herein (PDB: 6PXN), using chain A to simulate the 'loop up' conformation and chain B to simulate the 'loop down' conformation. As starting structures for the WT CK1δ simulations, we selected an apo structure (PDB: 1CKJ) of 2.46Å resolution, which has the activation loop crystallized both in 'up' (chain A) and 'down' (chain B) conformations (Longenecker et al., 1996). This structure contained two (chain A) or three (chain B) $WO_4^{-2}$ anions, which were computationally replaced by $SO_4^{2-}$ anions at the same positions, to make the simulations of WT CK1δ more comparable to the *tau* simulations. Using these structures, we created the systems to be simulated, described in Supplemental Table 5.

*Systems set up and equilibration:* The systems described in Supplemental Table 5 were refined by (i) position restrained energy minimization followed by (ii) full energy minimization, using Maestro (Schrödinger). The protonation states of the minimized models were estimated using the H++ server (Anandakrishnan et al., 2012) and hydrogens were added using pdb2pqr (Dolinsky et al., 2007).

All systems were solvated in a pre-equilibrated cubic TIP3P (Jorgensen et al., 1983) water box with at least 15 Å between the protein and the box boundaries. The net charge of the system was neutralized with $Na^+$ or $Cl^-$ counterions. Parameters for protein atoms and counterions were extracted from the ff14SB forcefield (Maier et al., 2015), while parameters for the $SO_4^{2-}$ anions were extracted from the Generalized Amber Force Field (GAFF) (Wang et al., 2004) and adjusted as proposed by (Kashefolgheta and Vila Verde, 2017).

Minimization and equilibration were performed with AMBER 16 (Case et al., 2016), using the following protocol: (i) 2000 steps of energy minimization with a 500 kcal $mol^{-1}$ $Å^{-1}$ position restraint on protein and $SO_4^{2-}$ anions; (ii) 1000 steps of energy minimization with a 500 kcal $mol^{-1}$ $Å^{-1}$ position restraint on protein atoms only; (iii) 2000 steps of energy minimization without position restraints; (iv) 50 ps of NVT simulation, with gradual heating to a final temperature of 300 K, with 10 kcal $mol^{-1}$ $Å^{-1}$ position restraint on protein and $SO_4^{2-}$ anions; (v) 1 ns of NPT simulation to equilibrate the density (or final volume of the simulation box).

**Conventional (cMD) and Gaussian Accelerated MD (GaMD) simulations**

Before running the GaMD simulations, we ran 100 ns of conventional MD simulations for each system, using AMBER 16 (Case et al., 2016) These simulations were performed in the NVT regime, with a time step of 2 fs, and all bonds involving hydrogen atoms were restrained with SHAKE (Rykckaertr et al., 1977). The PME method (Darden et al., 1993) was used to calculate electrostatic interaction using periodic boundary conditions, and a 12 Å cutoff was used to truncate non-bonded short-range interactions.

The final conformations produced by the cMD simulations were used as starting configurations for the GaMD simulations, which were performed with AMBER 17 (Case et al., 2017). For these simulations, additional acceleration parameters were used to boost the exploration of the conformational space, as described in (Miao et al., 2015). All systems had a threshold energy $E = V_{max}$ and were subjected to a dual boost acceleration of both the dihedral and the total potential energies. To optimize the acceleration parameters we first ran 2 ns of MD simulations with no boost potential, during which the minimum, maximum, average and standard deviation ($V_{min}$, $V_{max}$, $V_{av}$, $\sigma_{avg}$) of the total potential and dihedral energies were estimated and used to derive boost potentials as detailed in (Miao et al., 2015). These potentials were used to start 50 ns of Gaussian accelerated MD simulations, during which the boost statistics and boost potentials were updated until the maximum acceleration was achieved. The maximum acceleration was constrained setting the upper limit of the standard deviation of the total boost potential to be 6 kcal/mol.

We ran 5 replicas of production GaMD simulations for each system with fixed acceleration parameters derived from the previous equilibration stage. Each replica started from the same initial conformation, but the atoms were given different initial velocities, consistent with a Maxwell-Boltzmann distribution at 300 K. Each production simulation ran for 500 ns, totalizing 2.5 μs of sampling for each system, and 15 μs in total.

**Analysis of GaMD simulations**

*Conformational dynamics of loops:* The RMSD of the activation loop (Figure 4A-D) or loop L-EF (Figure 4E-H) was measured throughout the GaMD trajectories with CPPTRAJ using the *rms* module (Roe and Cheetham, 2013).

*Volumetric analysis of the substrate binding cleft:* The volume and shape of the substrate binding cleft and adjacent anion binding sites were obtained from the GaMD trajectories using

POVME 3.0 (Wagner et al., 2017). For each trajectory, conformations of WT or *tau* CK1δ were extracted every 2 ns, stripping all water molecules, counter-ions and sulfate anions. All protein conformations were superimposed to the same reference frame and the substrate binding cleft and adjacent anion binding sites were encompassed by three overlapping spheres as shown in Supplemental Figure 4G. POVME was then used to estimate the free (or empty) volume within the three overlapping spheres for all conformations extracted from the GaMD trajectories. The resulting volumes were averaged to produce three-dimensional density maps, as showed in Figure 4 (panels I-L). The maps in Figure 4 were contoured at 0.10 and represent regions that are found more frequently 'open' during the simulations.

*Principal Modes of Motion:* To detect the principal modes of motion displayed by WT and *tau* CK1δ, we used atomic fluctuations sampled during the GaMD simulations to perform Principal Component Analysis (PCA) (Amadei et al., 1993; Amadei et al., 1996). Before performing PCA, we stripped the trajectories of all solvent, ions and protein side-chains, keeping the backbone atoms only. We then concatenated and aligned these new trajectories to the same reference frame. To construct and diagonalize the co-variance matrix of atomic fluctuations, we used the *matrix* and *analyze* modules, respectively, in CPPTRAJ (Roe and Cheetham, 2013). For each system (WT CK1$\delta^{\text{loop down}}$, WT CK1$\delta^{\text{loop up}}$, *tau* CK1$\delta^{\text{loop down}}$, *tau* CK1$\delta^{\text{loop up}}$), we projected their respective trajectories into the obtained eigenvector space using the *projection* function in CPPTRAJ (Roe and Cheetham, 2013). The projections of each system along the subset of the eigenvector space formed by the 1$^{st}$ and 2$^{nd}$ principal components are shown in Figures 5 and S5.

All visualization of the GaMD simulations was performed with VMD (Humphrey et al., 1996).
